# PersonaDrive: A Method for the Identification and Prioritization of Personalized Cancer Drivers

**DOI:** 10.1101/2021.10.11.463919

**Authors:** Cesim Erten, Aissa Houdjedj, Hilal Kazan, Ahmed Amine Taleb Bahmed

## Abstract

**Motivation:** A major challenge in cancer genomics is to distinguish the driver mutations that are causally linked to cancer from passenger mutations that do not contribute to cancer development. The majority of existing methods provide a single driver gene list for the entire cohort of patients. However, since mutation profiles of patients from the same cancer type show a high degree of heterogeneity, a more ideal approach is to identify patient-specific drivers.

**Results:** We propose a novel method that integrates genomic data, biological pathways, and protein connectivity information for personalized identification of driver genes. The method is formulated on a personalized bipartite graph for each patient. Our approach provides a personalized ranking of the mutated genes of a patient based on the sum of weighted ‘pairwise pathway coverage’ scores across all the patients, where appropriate pairwise patient similarity scores are used as weights to normalize these coverage scores. We compare our method against three state-of-the-art patient-specific cancer gene prioritization methods. The comparisons are with respect to a novel evaluation method that takes into account the personalized nature of the problem. We show that our approach outperforms the existing alternatives for both the TCGA and the cell-line data. Additionally, we show that the KEGG/Reactome pathways enriched in our ranked genes and those that are enriched in cell lines’ reference sets overlap significantly when compared to the overlaps achieved by the rankings of the alternative methods. Our findings can provide valuable information towards the development of personalized treatments and therapies.

**Availability:** All the code and necessary datasets are available at https://github.com/abu-compbio/PersonaDrive.

## 1 Introduction

Cancer is an evolutionary and complex disease arising in many cases from the dysregulation of gene sequence, expression, genomic and transcriptomic alterations, in which abnormal cells divide uncontrollably and can invade nearby tissues (Masica and Karchin, 2011; Vogelstein *et al.*, 2013). The mutations that give a cancer cell a fundamental growth advantage and promote cancer development are called *driver mutations* and the corresponding genes subject to alteration are called *driver genes*. On the other hand there may exist several other mutations that occur randomly in a tumor sample but that are not directly associated with cancer development. Such mutations are the so called *passenger mutations*. A major challenge in cancer genomics is to distinguish the drivers that are causally linked to cancer from the passenger mutations (Haber and Settleman, 2007; Aburatani, 2011; Adjiri, 2017).

Due to the advances in high-throughput DNA sequencing technology, projects such as The Cancer Genome Atlas (TCGA) (Weinstein *et al.*, 2013) and The Cancer Cell Line Encyclopedia (CCLE) (Barretina *et al.*, 2012) have been able to systematically generate genomic data for thousands of tumors and cell lines across many cancer types providing useful data sources for many cancer driver identification studies. Several related versions of the cancer driver identification problem have been proposed. In one version the goal is to come up with a set of driver modules where each module consists of genes expected to be important in the development of the cancer under study (Leiserson *et al.*, 2015; Ciriello *et al.*, 2012a; Vandin *et al.*, 2012; Zhao *et al.*, 2012; Leiserson *et al.*, 2013; Babur *et al.*, 2015; Kim *et al.*, 2015; Ahmed *et al.*, 2019; Baali *et al.*, 2020). Another problem version sets the goal as that of ranking the altered genes with respect to their potentials for being cancer drivers. The methods suggested for this purpose can further be classified into two depending on whether the goal is to provide a cohort-level driver gene prioritization or a patient-specific driver gene prioritization.

The cohort-level driver gene identification and prioritization methods usually integrate multi-dimensional genomic data (Pham *et al.*, 2021; Cheng *et al.*, 2016). These methods can be categorized into different classes such as frequency-based, hotspot-based, and network-based methods (Han *et al.*, 2019). MutsigCV is one of the notable approaches among the first class. It investigates mutational heterogeneity patterns by correcting for variation using patient-specific mutation frequency and spectrum, and gene-specific background mutation rates incorporating expression level and replication time (Lawrence *et al.*, 2013). A representative of the methods of the second class is OncodriveCLUST which detects driver genes with a significant bias toward mutations clustered within specific protein sequence regions (Tamborero *et al.*, 2013). Network-based methods constitute a class of promising methods to prioritize low-frequency and high-frequency cancer genes due to their power to elucidate molecular mechanisms and their ability to model gene interactions (Wei *et al.*, 2021). DriverNet is among the notable approaches that employ mutation data, gene expression, and biological network data to prioritize mutated genes based on their degrees of network connectivity to the differentially expressed genes (DEGs) in tumor samples (Bashashati *et al.*, 2012). The bipartite graph model employed by DriverNet to relate the set of mutated genes to the set of DEGs has inspired many subsequent driver identification methods. BetweenNet is one such recent method which utilizes a measure based on betweenness centralities of genes in protein-protein interaction (PPI) networks constructed for each patient to identify dysregulated genes and employs a random-walk process on the network to prioritize driver genes (Erten *et al.*, 2021).

### 1.1 Patient-Specific Driver Ranking Methods

It is well-known that the mutation profiles of patients from the same cancer type show a high degree of heterogeneity. Since each patient may have a distinct set of driver genes, a more ideal approach is to identify patient-specific drivers. There are two main challenges associated with personalized driver gene rankings; some patients may carry too many known driver genes, necessitating narrowing down the driver genes to the true drivers for that patient while on the other hand, some patients do not possess any known drivers, so it is necessary to discover novel drivers for those patients (Dinstag and Shamir, 2020). There have been several attempts to address these problems and similar to some cohort-level network-based driver identification methods, many of the methods within the personalized setting of the problem are also inspired by the concept underlying the DriverNet approach which relates mutations to their consequent effects on transcription and gene expression by utilizing an underlying interaction network or pathway information. DawnRank prioritizes personalized driver genes by quantifying the impact of the mutated gene on the DEGs using a random walk process (Hou and Ma, 2014). The single-sample controller strategy (SCS) implements network control theory for personalized driver gene identification by searching for the minimal set of mutated genes to control the maximal coverage of individual DEGs in a directed biological interaction network (Guo *et al.*, 2018). Prodigy employs the expression and mutation profiles of the patient along with data on known pathways and the connectivity information to prioritize personalized cancer driver genes using the Steiner tree model based on the impact of mutated genes on dysregulated pathways which are significantly enriched in DEGs (Dinstag and Shamir, 2020).

We observe two shortcomings of the existing personalized driver prioritization methods. One stems from the way they employ the available data. Although each one employs its algorithm on a specific sample from some cohort, it does so by neglecting the availability of data pertaining to other samples in the cohort. This is a waste of valuable data, as data from other samples may guide the personalized driver rankings of a specific sample. The emphasis of the personalized setting of the problem should not concern the employed input data but rather solely the provided outputs so that the provided gene rankings should be patient-specific. Yet another important drawback is the lack of evaluation methodology consistent with the personalized setting of the problem. Mainly two different approaches are employed in evaluations comparing alternative personalized driver ranking methods. In the first one the relevant method is employed on each sample from an available cohort and then the quality of the output is determined by cohort-wise aggregation of the patient-specific outputs. Usually a Condorcet voting is employed for such an aggregation and the resulting gene set is compared against a set of known reference genes such as the COSMIC Cancer Gene Census (CGC) database (Tate *et al.*, 2019). DawnRank, SCS, and IMCDriver (Zhang *et al.*, 2021b) are among the methods following this approach. Such an evaluation based on *ranking-aggregation-evaluation (RAE)* is flawed for the obvious reason; the patient-specific findings are lost in cohort-wise aggregation and a cohort-based prioritization method may provide better results than a patient-specific driver ranking method. Another evaluation strategy employed by PRODIGY is to compare the rankings of each sample against the reference set of known drivers separately and then aggregate the results by averaging values for the entire cohort as a function of the top *N* ranked genes. If a sample has less than *N* ranked genes, the last value for that sample is duplicated so that all quality measure vectors for all patients are of length *N*. Although not as severe as the RAE strategy, this evaluation strategy based on *ranking-evaluation-aggregation (REA)* suffers from certain shortcomings as well. The first issue is that any set of reference known cancer drivers, although appropriate for cohort-based driver prioritization settings, is not an appropriate ultimate golden standard for the personalized setting of the problem. If used in such a setting, the evaluations must be supported by other strategies that can emphasize the personalized nature of the problem. Secondly, even if the issue with the golden standard is neglected, the duplication procedure of the REA strategy is problematic; a hypothetical prioritization providing a single gene such as TP53 as its output for every sample would superficially outperform the personalized driver identification and ranking methods.

### 1.2 Novelties of PersonaDrive

In this work, we propose a novel method called PersonaDrive that integrates genomic data, transcriptomic data, protein-protein interaction, and biological pathway data for the identification and prioritization of personalized driver genes. Unlike other network-based personalized driver identification and ranking methods PersonaDrive takes into account the data available from the whole cohort while producing the personalized ranking for every specific patient. A *pairwise patient similarity* score based on the amount of overlap between the set of DEGs of the pair is employed in determining the degree of the influence each patient’s data has on the other’s personalized driver ranking. Although the idea of exploiting data from the whole cohort exists in methods based on machine learning such as the IMCDriver, network-based methods not employing learning have thus far not incorporated the concept. Furthermore we note that a learning-based method like IMCDriver that employs the RAE strategy for the evaluations suffers even more than its nonlearning-based counterparts, since the reference set of known drivers is already part of the input features employed in learning. In fact the only other set of features employed in the learning stage of the method is simply the mutations data of the samples. This makes the prioritizations of such methods highly biased towards the reference set and evaluating such prioritizations based on the RAE strategy applied with the same set of reference genes may lead to incorrect conclusions. A second novelty of PersonaDrive is the concept of *pairwise pathway coverage* based on which the rankings of mutated genes are determined. Inspired by the original DriverNet model building on the influence of mutated genes on the DEGs and the subsequent methods DawnRank, PRODIGY, and SCS all of which employ a similar idea, we also create a network model of such influences. However different from these methods, PersonaDrive’s ranking score is based on the amount of coexistence of the mutated gene and the DEG pair in biological pathways, that is pairwise pathway coverage. Informally a mutated gene of a sample coexisting with many DEGs in many pathways gets a higher score and this score is magnified even more if it does so in many *similar* samples. Finally, another contribution of this work is with regard to the proposed evaluation framework. Notably we propose a modification to the REA strategy of PRODIGY to resolve the issues arising from the duplication idea. More importantly, we propose a novel framework for the evaluation of personalized driver identification and ranking methods. This framework is based on the idea of creating a separate reference golden standard set for each sample. Since the cell line data is usually richer than the tumor data available through TCGA in terms of personalized features, the framework is built on employing such data. A reference gene set is constructed for each cell line sample by incorporating the drug sensitivity, the drug targets data together with biological pathways data.

## 2 Methods

We first provide a description of the PersonaDrive method, a novel personalized cancer driver identification and ranking algorithm. Next, we provide a description of the introduced framework for the evaluation of cancer driver identification and prioritization outputs. The proposed evaluation framework includes certain novelties directed specifically towards the personalized setting of the problem.

### 2.1 PersonaDrive Algorithm

Figure 1 provides an overview of the PersonaDrive algorithm which consists of three main steps: i) constructing personalized bipartite networks; ii) computing edge weights in the bipartite networks; iii) personalized ranking of the genes. Below we provide a description of each of these steps in detail.

**Fig. 1.**
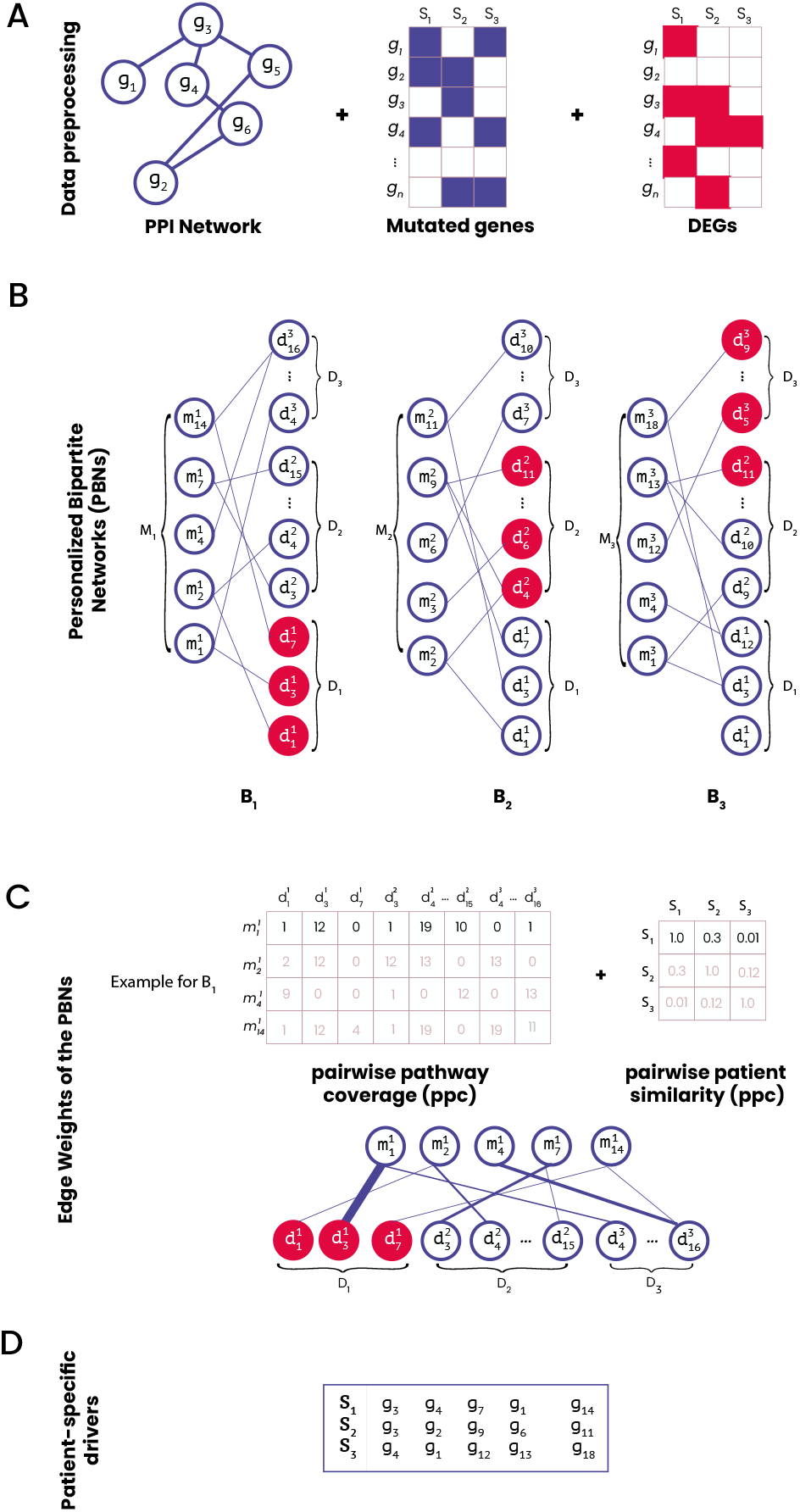
A depiction of the main steps of the PersonaDrive algorithm. A) Integrating the relevant genomic data (mutations and DEGs) with the interaction network. B) Constructing the personalized bipartite network *B*_*i*_ for each sample *S*_*i*_ in the cohort *S*. C) Computing the pairwise pathway coverage and the pairwise patient similarity values. The edge weights in each PBN *B*_*i*_ is then assigned based on these values D) The final output is a ranking of the mutated genes in *M*_*i*_ based on their influence scores which in turn are computed with respect to the weights of incident edges in the PBNs.

#### 2.1.1 Personalized Bipartite Networks (PBNs)

Similar to DriverNet, PersonaDrive constructs a bipartite graph to model the relationship between the set of mutated genes and DEGs. However different from DriverNet, PersonaDrive constructs a personalized network for each sample.

Let *G* = (*V, E*) denote the PPI network, where *V* denotes the set of nodes corresponding to the genes and *E* denotes the set of edges corresponding to pairwise protein-protein interactions. Assume the genes are denoted with *g*_1_, *g*_2_, … *g*_|*V*|_. Note that we use the same notation to denote a node of *G* and its corresponding gene. Let *P* = {*P*_1_, *P*_2_, … , *P*_*r*_} denote the set of functional pathways and *S* = {*S*_1_, … *S*_*n*_} denote the set of samples. If a gene *g*_*x*_ is mutated in sample *S*_*i*_ we create an instance of *g*_*x*_ denoted with 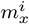 and denote the set of all such instances of genes mutated in sample *S*_*i*_ with *M*_*i*_. Similarly if a gene *g*_*y*_ is a DEG for sample *S*_*i*_, we create an instance of *g*_*y*_ denoted with 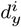 and denote the set of all such instances of DEGs of sample *S*_*i*_ with *D*_*i*_. For each sample, a gene is regarded as a DEG if its z-score > 1.0 or z-score < −1.0 based on its expression value in the sample as compared to its expression values in the whole population. Note that this is similar to the DriverNet’s *outlying gene* concept except that DriverNet employs a threshold of 2 rather than 1. Unlike the definitions employed by most personalized driver identification and ranking methods such as DawnRank, PRODIGY, and SCS, this definition of a DEG does not rely on the existence of paired normal/tumor data or background gene expression distributions from healthy samples of the same tissue of origin.

For each sample *S*_*i*_ we construct a bipartite graph *B*_*i*_ with the edge set *E*_*i*_. *B*_*i*_ has two node partitions, the first partition 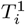 consists of nodes corresponding to the set of mutated genes *M*_*i*_. The second partition 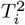 has nodes corresponding to the union of instances of DEGs of all the samples, that is *D*_1_ ∪ *D*_2_ ∪ … *D*_*n*_. Note that a gene may appear multiple times in the second partition as instances of different DEG sets from different samples. For 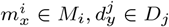, there exists an edge 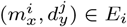, if (*g*_*x*_, *g*_*y*_) ∈ *E* and 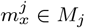. In other words, an instance of a gene *g*_*x*_ mutated in sample *S*_*i*_ has a bipartite edge with an instance of a gene *g*_*y*_ determined to be a DEG in sample *S*_*j*_, if they interact in the PPI network and *g*_*x*_ is a mutated gene in sample *S*_*j*_ as well. The subsequent steps assume all nodes in 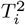 have degree greater than zero in *B*_*i*_. Therefore we remove all zero-degree nodes from the second partition.

#### 2.1.2 Edge Weights of the PBNs

The weight of an edge 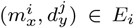 is computed with respect to a normalized value of the *pairwise pathway coverage (ppc)* score of incident nodes 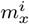 and 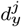. The score 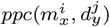 is defined as the number of pathways *P*_*k*_ ∈ *P* such that *g*_*x*_, *g*_*y*_ ∈ *P*_*k*_. The idea behind the *ppc* scoring is that the more the pathways shared by a mutated gene and DEG pair of a sample, the larger the potential for the mutated gene to be a cancer driver in that sample. Note that this is quite different from simply counting the number of pathways a mutated gene is involved in. For a mutated gene in a sample and every interacting partner that is a DEG in the sample, the pathways they coincide are counted towards the driver potential of the mutated gene in that sample. Furthermore since it is not necessarily the case that *i* = *j*, an edge weight is not computed in isolation, the data from a single sample being the sole determinant. It is also exposed to the mutation/dsyregulation patterns of the rest of the samples in the whole cohort, upto certain degrees formalized by the proposed normalization.

The normalization of the weight of an edge 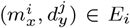 is with respect to the *pairwise patient similarity (pps)* value of the samples *S*_*i*_ and *S*_*j*_. Such a similarity is defined based on the overlap of the DEGs of the two patients. More specifically, for samples *S*_*i*_, *S*_*j*_ we define 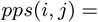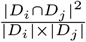. We note that similar overlap score definitions have been employed previously in measuring amount of overlaps among pairs of groups in various studies on clustering biological networks (Nepusz *et al.*, 2012; Baali *et al.*, 2020).

Incorporating the proposed normalization we define the edge weight of 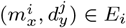 as,

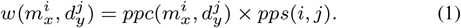

#### 2.1.3 Prioritizing mutated genes

For each mutated gene *g*_*x*_ in sample *S*_*i*_, we define an influence score 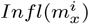 for the instance 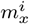 based on the weights of incident edges in the corresponding bipartite graph *B*_*i*_:

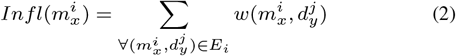

All mutated genes of *S*_*i*_ are prioritized with respect to this influence score and output in the relevant order by the PersonaDrive algorithm.

### 2.2 Comparative Evaluation Framework

We propose two main types of comparative evaluations emphasizing the personalized aspect of the problem. The first type of evaluations employs the reference sets used in previous work and a modified version of the REA strategy proposed in Dinstag and Shamir (2020). The second type of evaluations in line with the personalized nature of the problem proposes the use of novel personalized reference sets that are based on cell line data. We first construct reference sets by incorporating drug sensitivity data specific to each cell line sample and repeat the evaluation methodology with respect to the modified REA strategy. In order to complement the findings, we propose an additional evaluation method on cell line data that abandons the modified REA strategy as a way to quantify the matches between reference sets and output driver prioritizations but rather introduces a more flexible matching based on pathway enrichments.

#### 2.2.1 Evaluations with Reference Sets Relevant for Cohort Studies

The first type of evaluations is based on appropriate modifications of the REA strategy. Note that the REA strategy is employed by PRODIGY after producing a *personalized reference set* for each sample. The personalized reference set of a sample is considered to be the intersection of the set of mutated genes of the sample and an appropriate reference set of known cancer genes. The original strategy first fixes a certain desired output size *N*. In the case of PRODIGY evaluations *N* is set to 20. For every increment of *k* from 1 to *N*, for a benchmark method 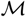 it compares the first *k* genes output by *M* in the ranking of each sample *S*_*i*_ against the personalized reference set of *S*_*i*_. It then aggregates the evaluation scores, usually measured in terms of precision and recall, by averaging the values over the entire cohort. If a method 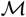 provides *k*′ ranked genes for a certain sample *S*_*i*_ for *k*′ < *N*, the evaluation score for *S*_*i*_ at instance *k*′ is repeatedly duplicated until *N* in the evaluation scores of 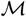.

We propose two main modifications to the original strategy. First of all, since not all samples are expected to have the same number of personalized cancer drivers, the first modification to this strategy consists of assigning *N* dynamically based on the sizes of the employed personalized reference sets. We define *N* to be twice the median of the sizes of the personalized reference sets after excluding the samples with reference set sizes less than three from the evaluations. Secondly due to the issues stemming from the duplication procedure of the original strategy, rather than duplicating the scores of 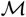 for *S*_*i*_ at instance *k*′ if *k*′ < *N*, we simply remove the sample *S*_*i*_ from all the successive average evaluation score calculations for the instances *k*′ + 1, … , *N*.

#### 2.2.2 Evaluations Based on Cell Line Data

The second type of evaluations are based on the observation that even the modified REA strategy may fail to provide conclusive findings if the personalized reference sets are constructed from known driver databases appropriate for population-based studies, such as the CGC (Tate *et al.*, 2019), NCG (Repana *et al.*, 2019), or CancerMine (Lever *et al.*, 2019). In order to resolve this issue and construct potential personalized reference sets based on data specific to each sample we propose the use of cell line data. For this type of evaluations, for each available cell line we define a novel reference gene set by compiling the target genes of drugs that are found to be sensitive based on data from GDSC and DepMap databases for that cell line. The reference set is further filtered with the CGC set of known drivers. Once the personalized reference sets are determined, the rest of the modified REA strategy described above is applied. The statistical information of the targets in GDSC and the DepMap reference sets for each cancer type under study are available in Supplementary Tables 1-2. The GDSC cell line drug sensitivity is retrieved from the GDSC2 dataset (Yang *et al.*, 2013). We label a cell line as *sensitive* to a drug if the z-score value is less than 0. DepMap cell line drug sensitivity data are gathered from the datasets of drugs screened with PRISM. Specifically, we used the ‘primary-screen-replicate-collapsed-log-fold-change’ of Corsello *et al.* (2020). The primary dataset consists of the results of pooled-cell line chemical-perturbation viability screens for 4518 compounds screened against 578 cell lines at a 2.5*μM* dose. We label a cell line as sensitive to a drug if the collapsed fold-change value is less than −0.8.

Finally, to extend the personalized emphasis proposed by the second type of evaluations employing the modified REA strategy together with the relevant cell line data we introduce one last type of evaluations again based on cell line data. The extension is based on the observation that directly comparing overlaps of sensitive drug target genes with the prioritized genes of a cell line can be too strict. Therefore we also evaluate the methods based on KEGG (Kanehisa *et al.*, 2019) and Reactome (Fabregat *et al.*, 2018) enrichment analysis by checking the amounts of overlaps between the pathways enriched significantly in the genes output by some personalized prioritization method and those that are enriched in cell line reference sets constructed from drug sensitivity data. In pathway analysis to avoid bias related to size it is suggested to employ gene sets of size at least 10 (Cirillo *et al.*, 2017). Therefore we filter out the samples with less than 20 genes in their reference sets. Furthermore, samples with less than 20 genes in the output prioritization of any method under comparison are also excluded from the analysis. We find the set of enriched KEGG or Reactome pathways using the g:GOSt tool (the core of g:Profiler tool) of Raudvere *et al.* (2019) which maps genes to known functional information sources and detects statistically significantly enriched terms. For each cell line sample *S*_*i*_, we identify 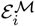, the set of pathways enriched significantly in the output prioritized gene set of a method 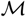 and 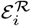 the set of pathways enriched significantly in the reference set of *S*_*i*_. The set of mutated genes *M*_*i*_ are employed as the background set in both cases. We then compute the *enriched pathway overlap (EPO)* score of a method 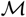 as the average of 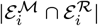 over all samples *S*_*i*_.

## 3 Results

We first compare PersonaDrive results against those of three existing personalized driver prioritization methods: Prodigy, SCS, and DawnRank. Our evaluations are based on patient data from TCGA project and cell line data from the CCLE project. Followed by the comparative evaluations we provide a more detailed analysis of the output prioritizations provided by the PersonaDrive algorithm.

### 3.1 Comparative Evaluations

All methods under comparison assume the same type of input data as PersonaDrive. Therefore no algorithm has any additional advantage of extra information. More specifically all the algorithms assume data from a cohort which include somatic mutation data and the gene expression data of the tumor samples, PPI network data, and pathway data. PRODIGY and PersonaDrive make explicit use of the pathway information, whereas DawnRank and SCS use it implicitly in the construction of the employed *gene network* which aggregates protein-protein interactions and pathway interactions. Actually, PersonaDrive is even less restricted than the other benchmark methods in terms of its input requirements in the sense that it requires the expression data only from the tumor samples. On the other hand Prodigy, SCS, and DawnRank additionally require gene expression data from the normal samples. We employed the TCGAbiolinks R package to compile the relevant data pertaining to the samples from two cancer types from TCGA (Colaprico *et al.*, 2016). The colon adenocarcinoma (COAD) cohort contains mutations and gene expression data of 396 tumor samples and the gene expression data of 41 normal samples, whereas the lung adenocarcinoma (LUAD) cohort contains mutations and gene expression data of 508 tumor samples and the gene expression data of 58 normal samples. We employed the DepMap database version 20Q1 (Dempster *et al.*, 2019; Ghandi *et al.*, 2019; Meyers *et al.*, 2017) to gather the mutations and gene expression data of the cancer cell line samples. Regarding the expression data of normal cell line samples we employ the normal human bronchial epithelial (NHBE) cell line from the Expression Atlas database of Petryszak *et al.* (2016) for LUAD and the CCD-18Co human normal colon myofibroblasts data from Ferrer-Mayorga *et al.* (2019) for COAD. For both the TCGA and the CCLE datasets, we filter out the silent mutations from the somatic mutation data.

We employ two different interaction networks in our evaluations; the STRING network employed in Dinstag and Shamir (2020) and the DawnRank gene interaction network of Hou and Ma (2014) which is the network employed in the SCS study as well. The former network consists of experimentally validated physical interactions with confidence score greater than 0.7. The resulting network contains 11,302 nodes and 273,210 edges. The latter is compiled from multiple sources including the network used in MEMo (Ciriello *et al.*, 2012b) as well as the curated information from the Reactome and KEGG pathway databases, and the NCI-Nature Curated PID (Schaefer *et al.*, 2009). It contains 11,648 nodes and 209,326 edges. The experimental results presented in the main document are those obtained with the latter network, whereas the results with the STRING network are provided in the Supplementary Document. Finally, the employed pathway data consist of the KEGG pathways retrieved from the supplementary material of Dinstag and Shamir (2020).

#### 3.1.1 Comparisons with Reference Sets Relevant for Cohort Studies

The personalized reference sets are constructed with respect to several relevant reference sets of known cancer genes. The first reference set employs Cancer Gene Census (CGC) and is denoted with *CGC*_*all*_. We further extract cancer type-specific known drivers from it by checking the ‘Tumor Types (Somatic)’ information. The resulting set is denoted with *CGC*_*specific*_. We also compile cancer type-specific genes from the Network of Cancer Genes (NCG) by filtering the *primary site* column. This reference set is denoted with *NCG*_*all*_. We further identify a subset of it as the intersection of *NCG*_*all*_ and *CGC*_*all*_. Hereafter, this reference gene set is referred to as *NCG*_*CGC*_. The third repository, CancerMine, uses text-mining to catalogue cancer associated genes where it also extracts information about the type of the cancer. We compile a list of genes that have at least two citations in literature and denote the resulting reference set with *CancerMine*_*all*_. The last reference set is constructed by intersecting *CancerMine*_*all*_ cancer-specific genes with the set of all CGC genes *CGC*_*all*_ and is denoted with *CancerMine*_*CGC*_ The number of genes in each reference is provided in Supplementary Table 3.

We evaluate each method based on how well it recovers the personalized reference set of each sample which is constructed by intersecting the set of mutated genes of the sample and one of the reference sets described above. The evaluation strategy is described in Section 2.2.1. Figure 2 shows the F1 score values for all the employed methods for the COAD and LUAD datasets from TCGA (see Supplementary Fig. S1 for the mean precision, and recall values). Here, the DawnRank network is employed as the input interaction network and the reference set used in the construction of the personalized references is *CGC*_*specific*_. We note that *N* is determined as 8 and 6 based on the sizes of the employed personalized reference sets for the COAD and LUAD datasets, respectively. For COAD, we observe that PersonaDrive achieves a higher performance than the alternatives in terms of all the metrics under consideration. It is followed by DawnRank, SCS, and Prodigy. PersonaDrive achieves the top performance for the LUAD dataset as well. It is followed by DawnRank, Prodigy, and SCS. When we construct each personalized reference set with respect to the *NCG*_*CGC*_ genes or *CancerMine*_*CGC*_ genes, we still observe that PersonaDrive performs better than its alternatives (Supplementary Fig. S2-S3). We evaluated the methods with three additional reference sets: *CGC*_*all*_, *NCG*_*all*_, and *CancerMine*_*all*_ (Supplementary Fig. S4-S5). PersonaDrive achieves the top performance for all these datasets and for all the metrics except for the LUAD cohort evaluations with respect to the reference sets constructed based on *CGC*_*all*_. For this experiment DawnRank achieves a slightly better F1 score than PersonaDrive. We also evaluate the sensitivity of our results to the employed input interaction network. To this end, we repeat the analogous experiments with the STRING network; see Supplementary Fig. S6-S9. The results show that PersonaDrive’s top performance do not depend on the employed interaction network. Note that SCS is not included for some evaluations on the STRING network as it does not execute until completion within a reasonable amount of time.

**Fig. 2.**
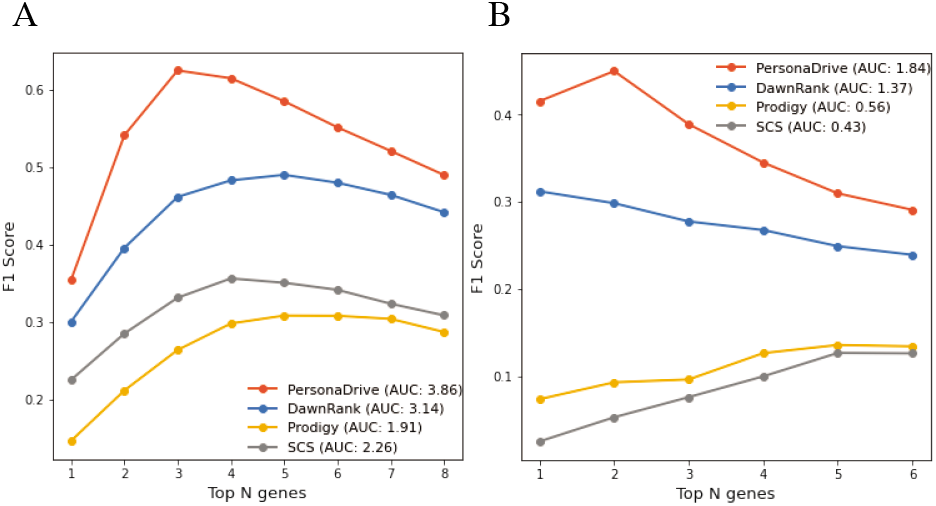
Comparison of the PersonaDrive outputs with those of the three alternative methods, DawnRank, SCS, and PRODIGY in terms of average F1 values for A) TCGA COAD dataset and B) TCGA LUAD dataset. DawnRank network is used as the input interaction network and *CGC*_*specific*_ is used as the reference set.

#### 3.1.2 Comparisons Based on Cell Line Data

Next, we compare the performances of the algorithms based on the personalized reference sets constructed with respect to the cell line data and evaluated with the modified REA strategy as described in Section 2.2.2. Figure 3 shows the mean F1 values for all the employed methods for the COAD and LUAD cell lines’ data where DawnRank network is used as input (see Supplementary Fig. S10 for the mean precision, and recall values). We note that *N* is determined as 34 and 10 for the COAD and LUAD datasets, respectively. PersonaDrive achieves the top performance in terms of all three metrics on the COAD dataset. DawnRank ranks second and SCS ranks third. Lastly, Prodigy shows the worst performance. Similarly, for the LUAD cell lines, PersonaDrive outperforms the alternatives. It is followed by DawnRank, Prodigy, and SCS in the order of decreasing performance. We repeat the analogous experiments with the STRING network; see Supplementary Fig. 11. The results show that PersonaDrive achieves a higher performance than the alternatives. It is followed by DawnRank and Prodigy.

**Fig. 3.**
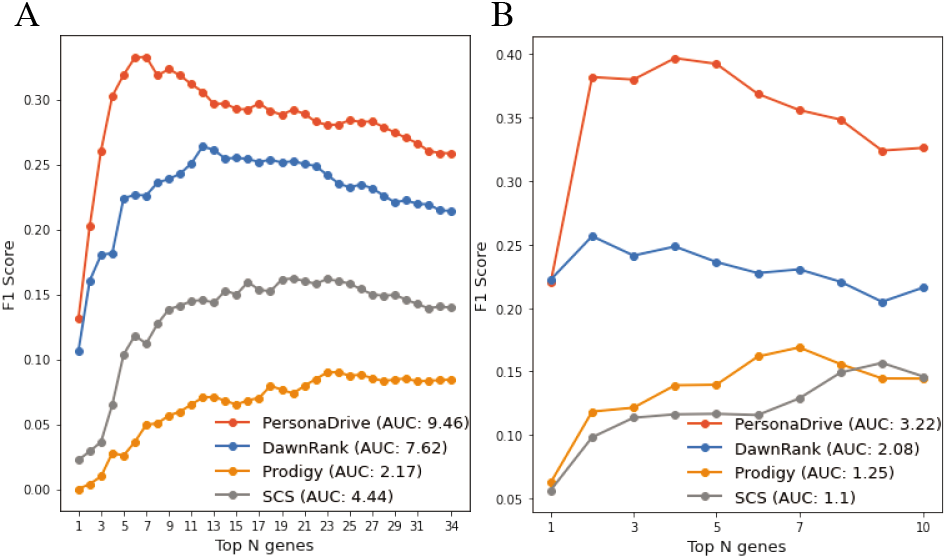
Comparison of the PersonaDrive outputs with those of the three alternative methods, DawnRank, SCS, and PRODIGY in terms of average F1 values for A) CCLE COAD cell lines and B) CCLE LUAD cell lines. DawnRank network is used as the input interaction network and the reference set is defined for each cell line based on the targets of sensitive drugs.

We perform an additional evaluation on the cell line datasets where we utilize pathway enrichment analysis as described in the second part of Section 2.2.2. This evaluation is performed only on the COAD data set since the set of pathways enriched significantly in cell line references constructed from drug sensitivity data is empty for 19 out of 27 LUAD cell lines under consideration. For COAD dataset, PersonaDrive performs the best in terms of *EPO* scores. Its *EPO* scores are 23.73 and 9.46 for KEGG and Reactome pathways, respectively. The second best performer is DawnRank with *EPO* scores of 12.27 and 8.15, respectively for the KEGG and Reactome pathways. The analogous scores of PRODIGY and SCS are 1.87, 1.62 and 0.73, 0.77, respectively. Results with the STRING network are similar and are available in Supplementary Table 4. Additionally, we investigate the most commonly enriched KEGG pathways across the cell lines with respect to their reference sets and determine whether these pathways are also found enriched within the sets of genes output by the methods under consideration. The top 5 most commonly enriched KEGG pathways in cell line reference sets for COAD are shown in Table 1. We observe that PersonaDrive’s top 20 genes show enrichment for *Pathways in cancer*, *MAPK signaling pathways*, and *EGFR tyrosine kinase inhibitor resistance* in all the cell lines for which these pathways are found to be enriched in the corresponding reference sets. We note that MAPK and EGFR signalling pathways are closely linked to colon cancer development (Fang and Richardson, 2005; Krasinskas, 2011). For the *Calcium signaling pathway* and *Central carbon metabolism* pathways, PersonaDrive achieves the largest overlap as well. Analogous results based on Reactome pathways can be found in the Supplementary Document.

**Table 1.** The top 5 most commonly enriched KEGG pathways in cell line reference sets and the corresponding numbers for the outputs of the methods under consideration.

|  | # of cell lines | PersonaDrive | DawnRank | Prodigy | SCS |
| --- | --- | --- | --- | --- | --- |
| *Pathways in cancer* | 11 | 11 | 11 | 3 | 7 |
| *MAPK signaling pathway* | 11 | 11 | 2 | 1 | 0 |
| *Calcium signaling pathway* | 10 | 8 | 0 | 0 | 0 |
| *Central carbon metabolism in cancer* | 9 | 7 | 1 | 0 | 0 |
| *EGFR tyrosine kinase inhibitor resistance* | 9 | 9 | 7 | 1 | 0 |

### 3.2 Further Analysis of PersonaDrive Gene Rankings

We provide an in-depth analysis of the prioritizations provided by the PersonaDrive algorithm. For this purpose we first assess whether PersonaDrive can identify rare drivers. We employ the TCGA dataset for this assessment since the number of samples in the cell line dataset is not appropriate for an analysis on rare drivers. We investigate the top *k* genes ranked by PersonaDrive for 1 ≤ *k* ≤ 20 and count the number of genes that belong to the following bins defined by the mutation frequencies and the CGC reference set of known drivers: < 2%(*CGC*_*all*_), 2 − 5%(*CGC*_*all*_),> 5%(*CGC*_*all*_), < 2%, 2 − 5%,> 5%. The first three bins correspond to the genes that belong to the set *CGC*_*all*_ and each of the rest of the bins contains genes not in *CGC*_*all*_. There is an additional *N/A* category since we predict less than 20 genes for some patients in the LUAD cohort. For both datasets, the top gene appears in the > 5%(*CGC*_*all*_) category most frequently. As *N* increases from 1 to 20 we see a decrease in the size of this category, whereas the sizes of the categories for rare known drivers and the categories for unknown drivers increase. Interestingly, for the LUAD dataset, we observe more genes in rare categories (both in *CGC*_*all*_ and not in *CGC*_*all*_) compared to the results obtained from the COAD dataset. Overall, these results show that PersonaDrive is able to prioritize rarely mutated genes and detect both rare and frequent known drivers (Supplementary Fig. S12).

Secondly, employing cell line data we implement two complementary types of analysis investigating the novel genes provided by PersonaDrive. In the first one the focus is on the distinctive genes provided for each of the cell lines employed in the evaluations, whereas in the second one the focus is on the common genes provided for most of the cell lines. With regard to the first type of analysis we relate the personalized drivers identified and ranked by PersonaDrive with the gene *essentiality* scores of DepMap computed based on systematic loss of function screens on cell lines (Meyers *et al.*, 2017). More specifically, for each sample we analyze the top ranked genes provided by PersonaDrive and check whether the novel ones among them, that is those not in the *CGC_all_* reference set of known drivers are verified for their driver potential by an external source such as DepMap essentiality. In order to emphasize the personalized nature of the setting of the problem under consideration, rather than simply using the essentiality scores we make use of the *preferentially essential genes* of a cell line as provided by DepMap. Note that a similar type of evaluation has been performed for evaluating novel cancer driver genes identified for a cohort of patients (Schulte-Sasse *et al.*, 2021). For a cell line and a gene, the *mean-subtracted score* is calculated by subtracting the mean score for that gene across all cell lines from the score of the gene in that cell line. Those with the lowest mean-subtracted scores are defined as the preferentially essential genes of the cell line. For each cell line, we compute the intersection between our top 20 ranked genes that do not appear in *CGC_all_* and the top 500 preferentially essential genes determined for that cell line. For LUAD, this intersection includes the genes *SOS*1 (ACH-000787), *POLR*2*B* (ACH-000587), *PTK*2 (ACH-000861), and *PLK*1 (ACH-000681), where the corresponding cell lines are indicated in parantheses. Among these genes, there is experimental evidence for SOS1’s oncogenic activity in lung cancer (Cai, 2019). PLK1 belongs to a family of serine/threonine kinases that are involved in cell-cycle regulation. PLK1’s overexpression is linked to tumor development for a variety of cancer types including non-small cell lung cancer (NSCLC) and its inhibitors have been designed for therapeutic purposes (Strebhardt and Ullrich, 2006). Similarly, PTK2 is a member of the non-receptor tyrosine kinase family and regultes cell survival, proliferation, migration and invasion (Aboubakar Nana *et al.*, 2019). Its inhibition has been explored as a potential therapy in both small cell and non-small cell lung cancer (Tong *et al.*, 2019). For COAD, the corresponding intersection includes the genes *HK*3 (ACH-001458), *GSK*3*B* (ACH-000957), *PTK*2 (ACH-000943). The complete list can be found in Supplementary Table 5. Among these genes, GSK3 inhibitors (GSK3i) have shown a strong synergistic effect with PARP inhibitors in a panel of colorectal cancer (CRC) cell lines with diverse genetic backgrounds (Zhang *et al.*, 2021a). HK3’s overexpression has been found to be associated with epithelial-mesenchymal transition in colorectal cancer (Pudova *et al.*, 2018). Additionally, overexpression of PTK2 has been correlated with metastatic colon cancer (Lark *et al.*, 2003; Miyazaki *et al.*, 2003; Tai *et al.*, 2016).

With regard to the second type of analysis, we perform a batch study on the novel genes proposed by PersonaDrive. For this, we take PersonaDrive’s most frequently prioritized genes among the top 20 ranking genes across all cell lines and explore the literature for associations with cancer. For lung cancer, Fibronectin 1 (FN1) which is ranked among the top 20 for four cell lines has been found to play critical roles in driving lung cancer (Spada *et al.*, 2021). For colon cancer, a recent study showed that ADCY9 which appears among the top 20 genes for five cell lines is playing a critical role and acts as an oncogene during colon cancer development (Yi *et al.*, 2018). The Insulin-like growth factor 1 receptor (IGF1R) is also supported by the recent studies as a candidate tumor suppressor and as an oncogene in colorectal cancer (Su *et al.*, 2014; Liu *et al.*, 2018). IRS2 has been shown to involve in tumorigenesis and is a candidate driver that is frequently upregulated in cancer development (Yun *et al.*, 2020).

## 4 Conclusion

Several crucial design choices are made in the PersonaDrive algorithm such as keeping the bipartite network and the set of pathways static throughout the ranking procedure, the z-score threshold used to determine the set of dysregulated genes, the calculation of pairwise patient similarity (pps) scores. We discuss each such choice and compare against the plausible alternatives in the Supplementary Document Section 3. Finally, to verify that the achieved evaluation results presented in the main document are not artifacts of the choice of the datasets and the specific evaluation strategies employed, we repeated our precision, recall, and F1 score evaluations on the same TCGA COAD data set as that employed in the most recent benchmark study, PRODIGY (Dinstag and Shamir, 2020). Furthermore for these evaluations we employed the unmodified REA strategy which is the original evaluation strategy proposed in the same study. We show that our conclusions remain almost the same in this controlled setting as well; see Supplementary Document Section 4.

To summarize, we provide a novel personalized driver identification and prioritization method, PersonaDrive. It relies on the simple yet powerful notions of *pairwise pathway coverage* and the *pairwise patient similarity*. Through extensive comparative evaluations we show that PersonaDrive outperforms three of the state-of-the-art personalized driver ranking methods. A major difficulty in the personalized setting of the driver gene ranking problem as compared to the impersonalized cohort setting has been the lack of appropriate evaluation methods and golden standard data reflecting the personalized nature of the problem. We propose several evaluation methods for this purpose; the modified REA strategy, its use with the personalized reference sets incorporating cell lines and drug targets data, and the pathway enrichment-based evaluations replacing the modified REA strategy are all novel contributions toward evaluations within the personalized setting of the problem.

One future direction of research is to extend our analyses to additional cancer types as experimental data on a larger set of cell lines become available. Another future direction is to utilize recent clinical datasets that include drug response data where several drugs are administered to different subsets of patients. In this way, we can define personalized reference sets for patients as well, by using a similar procedure to produce personalized reference sets for the cell lines.

## Acknowledgements

The authors are listed in the alphabetical order of their lastnames. We thank Gal Dinstag for providing details regarding the PRODIGY method.

