## Supplementary material for "PersonaDrive: A Method for the Identification and Prioritization of Personalized Cancer Drivers": PersonaDrive_supplementary_material.pdf

#### 1 Data Pre-processing

##### 1.1 Expression and Somatic Mutation data

We employed the TCGAbiolinks R package to compile the relevant data pertaining to the samples from two cancer types from TCGA [1]. The colon adenocarcinoma (COAD) cohort contains mutations and gene expression data of 396 tumor samples and the gene expression data of 41 normal samples, whereas the lung adenocarcinoma (LUAD) cohort contains mutations and gene expression data of 508 tumor samples and the gene expression data of 58 normal samples.. The gene expression values in the original dataset are in terms of $FPKM$ values, as an input to our method, we convert the $FPKM$ values to $TPM$ values as follows,

$$TPM_i = \frac{FPKM_i}{\sum(FPKM_j)} * 10^6 \tag{1}$$

However, read counts values for both normal and tumor samples are used as input for PRODIGY to perform differential expression analysis using DeSEQ2. Among the evaluated methods, only SCS requires expression data from paired normal and tumor samples. To evaluate all the employed methods on a larger dataset, we extract differentially expressed genes for SCS from unpaired gene expression data by retrieving the log-fold change values using DeSEQ2 R package.

For CCLE dataset, we pre-process both normal and tumor data matrices to contain only genes $G_{T,N}$ that exist in both the tumor and normal data as follows: $G_{T,N} = G_T \cap G_N$.

##### 1.2 Protein-protein Interaction Networks

We employ two different interaction networks in our evaluations; the STRING network employed in [2] and the Dawnrank gene interaction network of [3] which is the network employed in the SCS study as well. Prodigy employs the confidence values from the PPI network in its framework. We use the unweighted DawnRank network as input to PRODIGY by assigning edge weights to be 0.2 to all edges.

#### 2 Evaluation Metrics

The performance of each method is measured by computing the mean precision, recall, and F1 scores with respect to the reference sets. Let $D_i^{\mathcal{M}}$ be the set of $k$ genes output by $\mathcal{M}$ in the ranking of a sample $S_i$, and $R_i$ denotes the personalized reference of a sample $S_i$, then,

$$precision_i^{\mathcal{M}}[k] = \frac{|D_i^{\mathcal{M}}[k] \cap R_i|}{k} \tag{2}$$

$$recall_i^{\mathcal{M}}[k] = \frac{|D_i^{\mathcal{M}} \cap R_i|}{|R_i|} \tag{3}$$

$$F1_i^{\mathcal{M}}[k] = 2 * \frac{precision_i^{\mathcal{M}}[k] * recall_i^{\mathcal{M}}[k]}{precision_i^{\mathcal{M}}[k] + recall_i^{\mathcal{M}}[k]} \tag{4}$$

### 3 Design Choices of PersonaDrive

Several crucial design choices are made in the PersonaDrive algorithm. Each such choice needs to be discussed and compared against the plausible alternatives. Once the PBN of a patient is constructed and the influence scores are assigned to the nodes corresponding to the mutated genes of the patient, the strategy of PersonaDrive in ranking the driver genes is to *statically* rank them with respect to their scores, that is no updates on the employed data sets (PBN or the pathway data) are committed at any step throughout the ranking procedure. An alternative is to employ a *dynamic* ranking procedure which commits an update on the relevant structure(s) after each selection step. One possible type of dynamic updates is through the updates of the PBN. Once the initial scores are determined the largest scoring node is selected and assigned the best rank, and the PBN is updated by removing the selected node, its DEG neighbors in the PBN, and all the edges incident on these DEGs. Upon such an update the influence scores of all the mutated genes of the patient are recomputed. The procedure of selection made according to the new scores followed by the relevant updates is repeated until no DEGs remain in the PBN. Note that applying dynamic updates on relevant structures is the strategy followed by the DriverNet algorithm as well [4]. The version of PersonaDrive with relevant modifications is called PersonaDrive$_{dyn_nodes}$. Furthermore the overall design of PersonaDrive gives rise to a second possible dynamization - one based on dynamically updating the other relevant type of data available, that is the pathway data. For this second dynamization version which we call PersonaDrive$_{dyn_pathways}$, we keep track of a set of *uncovered* pathways for each sample individually. For each sample, we assign this set to the set of all KEGG pathways initially. Once the mutated gene with the largest influence score is selected, we identify the set of pathways that this gene and each of its connected DEGs coexist. We remove these pathways from the set of pathways that we have for this sample, since all such pathways are now 'covered' by at least one mutated gene-DEG pair of the sample. We repeat the evaluations based on reference gene sets with these two alternative dynamic versions of the PersonaDrive algorithm; see Supplementary Figures S13-S22. We observe that the original version of PersonaDrive performs significantly better than both of its dynamic versions in most of the evaluations. Exceptions are the results on TCGA COAD dataset, when Dawnrank or STRING network is used as input PPIs and the results on TCGA LUAD dataset when the reference set is $NCG_{all}$ and DawnRank network is used as input. More importantly, we observe a significant improvement of PersonaDrive over its dynamic versions for the CCLE datasets where we aim to define the reference sets in a more sample-specific manner than the reference sets appropriate for cohort-level settings of the driver identification problem.

A second design choice is with respect to the z-score threshold employed in determining whether a gene is considered a DEG. The analogous threshold employed in the DriverNet study is 2.0, whereas in PersonaDrive we employ a threshold of 1.0. This design choice stems from the fact that since DEGs are defined in a deterministic manner, employing a strict threshold such as 2.0, some less extreme but nonetheless important changes in expression that are modulated by a genomic event may be missed. On the other hand, employing a more flexible threshold such as 1.0 may potentially introduce false positive DEGs and thus may mislead the driver identification. However we note that the *ppc* concept of PersonaDrive, among its other roles, serves also as a mechanism to filter out such false positives as well. Nevertheless we implement another version of PersonaDrive and call it PersonaDrive $_{DEG\_threshold2}$, and compare the evaluation results of this version and the original PersonaDrive in Supplementary Figures S23-28. We observe that the original PersonaDrive performs better in all cases.

A third design choice of PersonaDrive is regarding the pairwise patient similarity (pps) definition, where the similarity of a pair of samples is defined based on a notion of similarity between the sets of DEGs of the samples. Instead of using only DEGs for the *pps* calculations an alternative is to merge the set of mutated genes and DEGs, and use the same notion of similarity between such pairs of sets to assign *pps* values. We call this version PersonaDrive$_{alt_similarity}$ and repeat the evaluations with this modified version; see Supplementary Figures S13-S22. We observe that PersonaDrive$_{alt_similarity}$ performs the same or worse than PersonaDrive in the majority of the cases for the relevant TCGA datasets. For the CCLE datasets, we observe that PersonaDrive performs better than PersonaDrive$_{alt_similarity}$ for LUAD samples when DawnRank network is used, whereas the opposite is observed for the COAD dataset. On the other hand, PersonaDrive$_{alt_similarity}$ shows a better performance than PersonaDrive when the STRING network is used for both CCLE datasets.

### 4 Comparisons on Original Dataset of PRODIGY

Finally, to verify that the achieved evaluation results are not artifacts of the choice of the datasets and the specific evaluation strategies employed, we repeated our precision, recall, and F1 score evaluations on the same TCGA COAD data set as that employed in the most recent benchmark study, PRODIGY. The reference set of known drivers, that is the CGC genes are also retrieved from PRODIGY. Note that in all the previous evaluations these two types of data are retrieved from the original sources of data and may be different from that employed by PRODIGY. Furthermore for these evaluations we employed the unmodified REA strategy which is the original evaluation strategy proposed in [2]. Note that with this strategy if a sample has less than $N$ ranked genes, the last value for that sample is duplicated so that all quality measure vectors for all patients are of length $N$. We demonstrate in Supplementary Figures S29-S31 that our approach outperforms the existing driver gene prioritization methods with a large margin under this controlled setting as well.

| | COAD | | | LUAD | | |
|---|---|---|---|---|---|---|
| | GDSC | DepMap | GDSC U DepMap | GDSC | DepMap | GDSC U DepMap |
| # of cell lines | 24 | 20 | 26 | 22 | 24 | 27 |
| # of empty reference sets | 7 | 0 | 2 | 10 | 0 | 2 |
| # of reference sets with size ≥ 3 | 10 | 15 | 17 | 1 | 16 | 16 |
| The median of reference set sizes | 9.5 | 16 | 17 | 3 | 5 | 5 |

Supplementary Table 1: The distribution of the number of cell lines in each dataset and some statistical information about cell line reference sets DawnRank $M_i$ is filtered by the set of genes in STRING network. The reference set for each cell line $i$ is defined as follows: $R_i = DrugTargets_i \cap M_i \cap CGC$.

| | COAD | | | LUAD | | |
|---|---|---|---|---|---|---|
| | GDSC | DepMap | GDSC U DepMap | GDSC | DepMap | GDSC U DepMap |
| # of cell lines | 24 | 20 | 26 | 22 | 24 | 27 |
| # of empty reference sets | 6 | 0 | 2 | 11 | 0 | 3 |
| # of reference sets with size ≥ 3 | 10 | 16 | 18 | 1 | 16 | 16 |
| The median of reference set sizes | 10.5 | 15 | 15 | 3 | 4.5 | 5 |

Supplementary Table 2: The distribution of the number of cell lines in each dataset and some statistical information about cell line reference sets where $M_i$ is filtered by the set of genes in STRING network. The reference set for each cell line $i$ is defined as follows: $R_i = DrugTargets_i \cap M_i \cap CGC$.

| | CGC | $CGC_{specific}$ | $NCG_{all}$ | $NCG_{CGC}$ | $CancerMine_{all}$ | $CancerMine_{CGC}$ |
|---|---|---|---|---|---|---|
| COAD | 723 | 61 | 156 | 43 | 121 | 45 |
| LUNG | 723 | 29 | 203 | 81 | 279 | 95 |

Supplementary Table 3: Size of reference sets for TCGA dataset

| | | KEGG | Reactome |
|---|---|---|---|
| COAD | PersonaDrive | **25.64** | **8.23** |
| | DawnRank | 7.07 | 4.85 |
| | Prodigy | 1.57 | 3.08 |

Supplementary Table 4: *EPO* scores of the methods under consideration for the KEGG and Reactome pathway enrichment-based evaluations on cell line data where STRING network is used as input.

| **Cell Line** | **Preferentially Essential Genes** | **Mutation Freq.** |
|---|---|---|
| | **LUAD** | |
| ACH-000787 | SOS1 | 4 |
| ACH-000587 | POLR2B | 6 |
| ACH-000861 | PTK2 | 4 |
| ACH-000681 | PLK1 | 8 |
| | **COAD** | |
| ACH-000501 | TLN1 | 9 |
| ACH-000957 | GSK3B | 3 |
| ACH-000943 | PTK2 | 4 |
| ACH-000202 | NUP155 | 7 |
| ACH-001458 | HK3 | 4 |
| ACH-000958 | CDC42 | 1 |
| ACH-001461 | TNNT2 | 3 |
| | PCF11 | 7 |
| ACH-000680 | PIP5K1B | 5 |

Supplementary Table 5: Preferentially essential genes identified by PersonaDrive for CCLE dataset where DawnRank network is used as input.

| | # of cell lines | PersonaDrive | DawnRank | Prodigy | SCS |
|---|---|---|---|---|---|
| Disease | 11 | 11 | 9 | 0 | 1 |
| Signal Transduction | 10 | 10 | 10 | 4 | 7 |
| Diseases of signal transduction by growth factor receptors and second messengers | 10 | 10 | 8 | 0 | 0 |
| Nervous system development | 8 | 3 | 0 | 2 | 0 |
| Axon guidance | 8 | 3 | 0 | 2 | 0 |

Supplementary Table 6: Top 5 Reactome pathways enriched by the most cell lines and the corresponding numbers for the outputs of the methods under consideration.

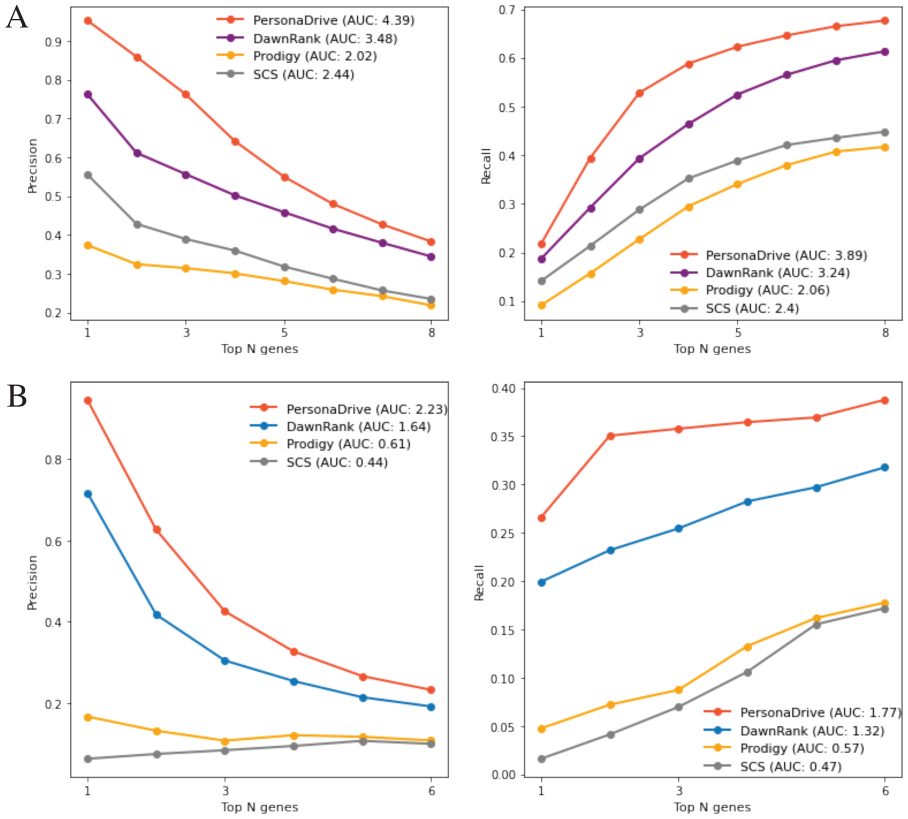


Supplementary Figure 1: Comparison of the PersonaDrive outputs with those of the three alternative methods, DawnRank, SCS, and PRODIGY in terms of average precision, and recall values for A) TCGA COAD dataset and B) TCGA LUAD dataset. DawnRank network is used as the input interaction network and $CGC_{specific}$ is used as the reference set.

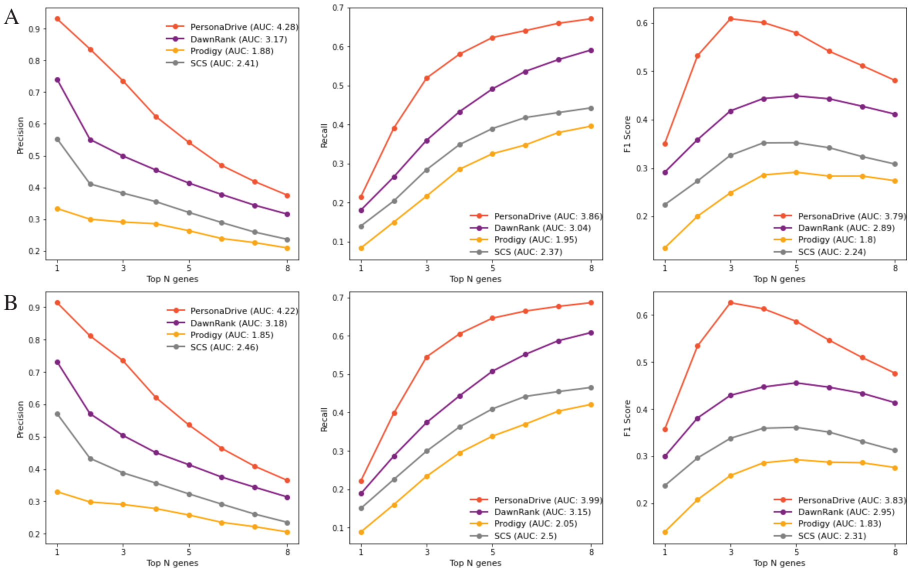


Supplementary Figure 2: Comparison of the PersonaDrive outputs with those of the three alternative methods, DawnRank, SCS, and PRODIGY in terms of average precision, recall, and F1 values for TCGA COAD dataset where DawnRank network is used as the input interaction network. A) $NCG_{CGC}$ genes are used as reference. B) $CancerMine_{CGC}$ genes are used as reference.

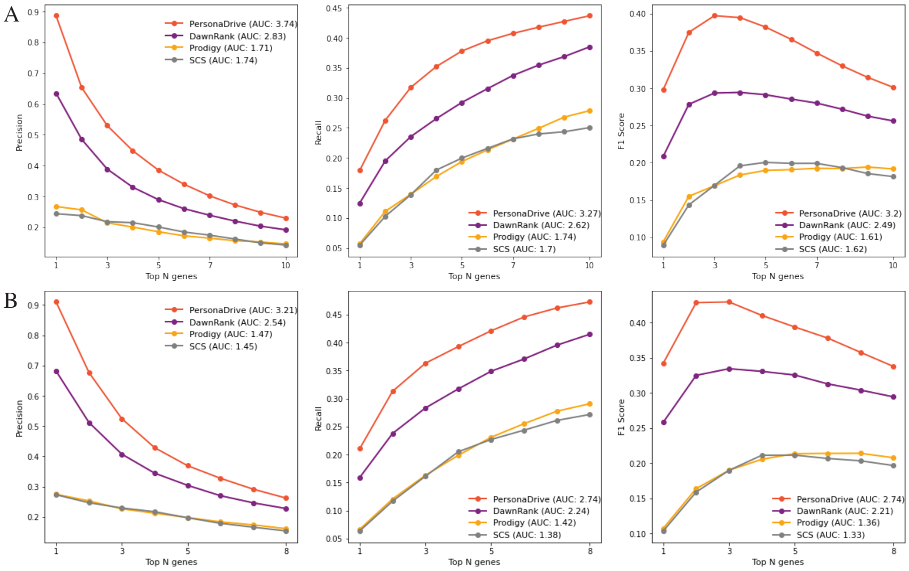


Supplementary Figure 3: Comparison of the PersonaDrive outputs with those of the three alternative methods, DawnRank, SCS, and PRODIGY in terms of average precision, recall, and F1 values for TCGA LUAD dataset where DawnRank network is used as the input interaction network. A) $NCG_{CGC}$ genes are used as reference set. B) $CancerMine_{CGC}$ genes are used as reference set.

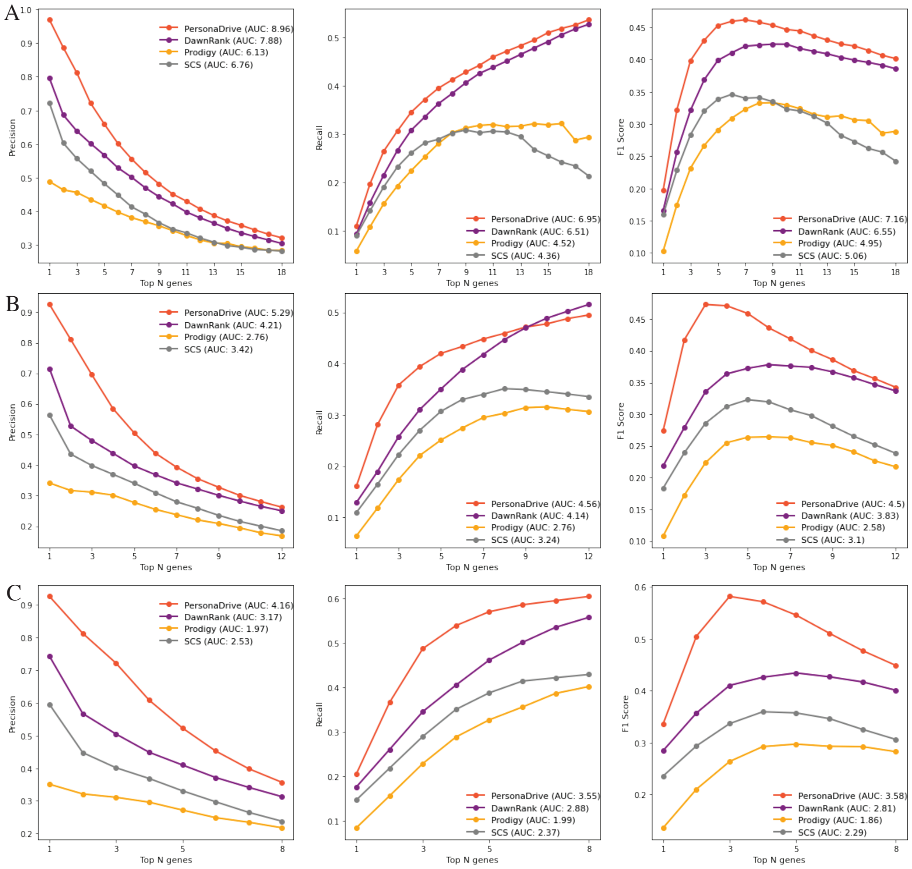


Supplementary Figure 4: Comparison of PersonaDrive with three alternative methods in terms of average precision, recall, and F1 values for TCGA COAD dataset where DawnRank network is used as the input interaction network. A) $CGC_{all}$ genes are used as reference. B) $NCG_{all}$ genes are used as reference set. C) $CancerMine_{all}$ genes are used as reference set.

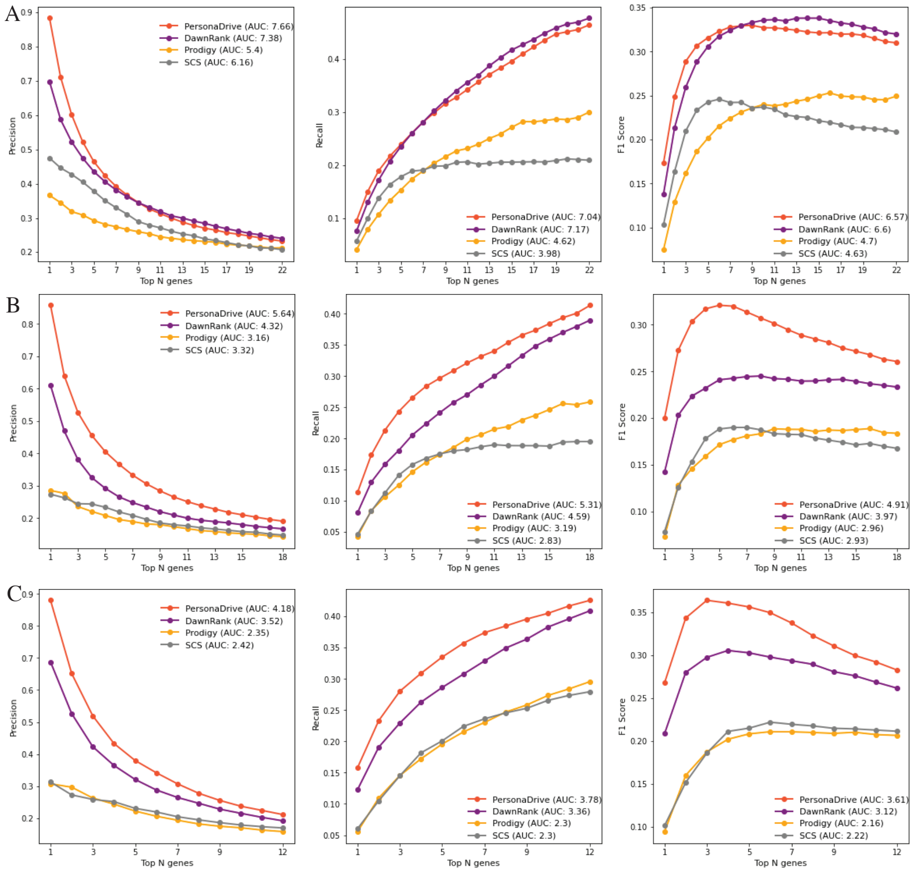


Supplementary Figure 5: Comparison of the PersonaDrive outputs with those of the three alternative methods, DawnRank, SCS, and PRODIGY in terms of average precision, recall, and F1 values for TCGA LUAD dataset where DawnRank network is used as the input interaction network. A) $CGC_{all}$ genes are used as reference set. B) $NCG_{all}$ genes are used as reference set. C) $CancerMine_{all}$ genes are used as reference set.

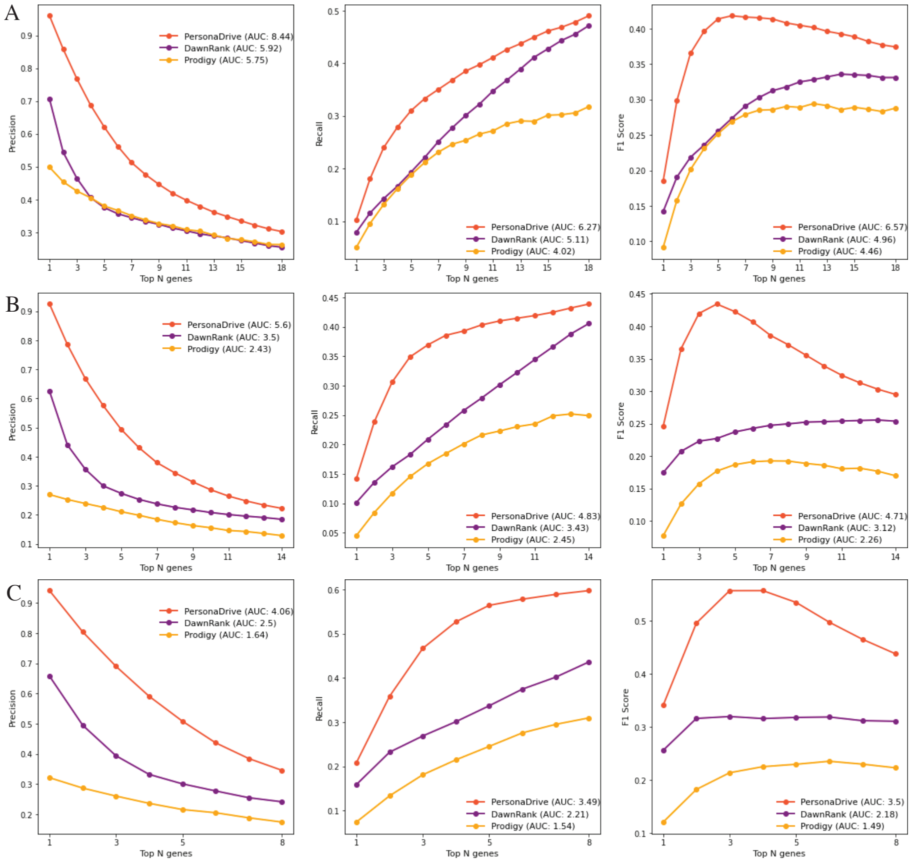


Supplementary Figure 6: Comparison of the PersonaDrive outputs with those of the three alternative methods, DawnRank, SCS, and PRODIGY in terms of average precision, recall, and F1 values for TCGA COAD dataset where STRING network is used as the input interaction network. A) $CGC_{all}$ genes are used as reference set. B) $NCG_{all}$ genes are used as reference set. C) $CancerMine_{all}$ genes are used as reference set.

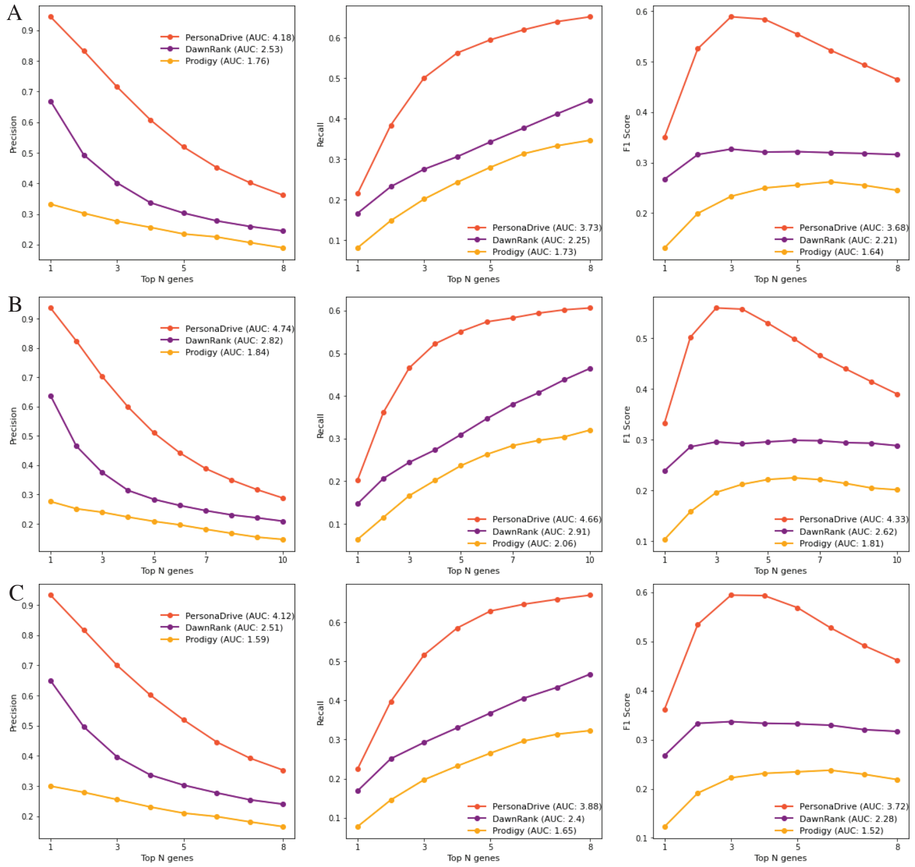


Supplementary Figure 7: Comparison of the PersonaDrive outputs with those of the three alternative methods, DawnRank, SCS, and PRODIGY in terms of average precision, recall, and F1 values for TCGA COAD dataset where STRING network is used as the input interaction network. A) $CGC_{specific}$ genes are used as reference set. B) $NCG_{CGC}$ genes are used as reference set. C) $CancerMine_{CGC}$ genes are used as reference set.

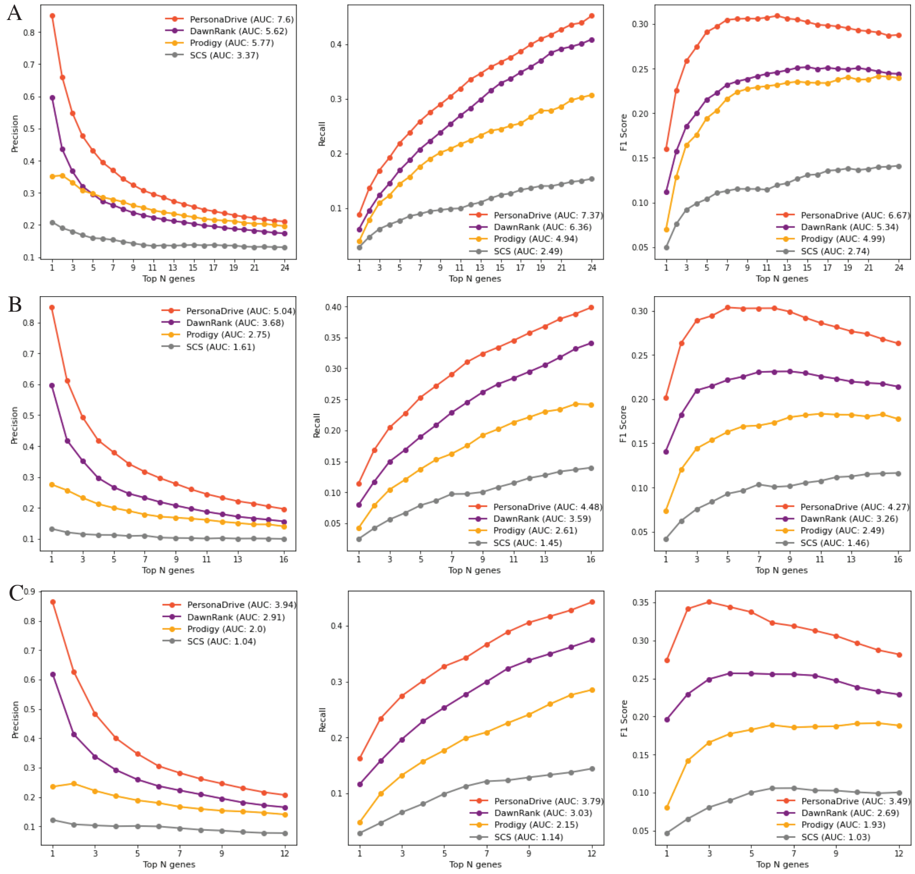


Supplementary Figure 8: Comparison of the PersonaDrive outputs with those of the three alternative methods, DawnRank, SCS, and PRODIGY in terms of average precision, recall, and F1 values for TCGA LUAD dataset where STRING network is used as the input interaction network. A) $CGC_{all}$ genes are used as reference set. B) $NCG_{all}$ genes are used as reference set. C) $CancerMine_{all}$ genes are used as reference set.

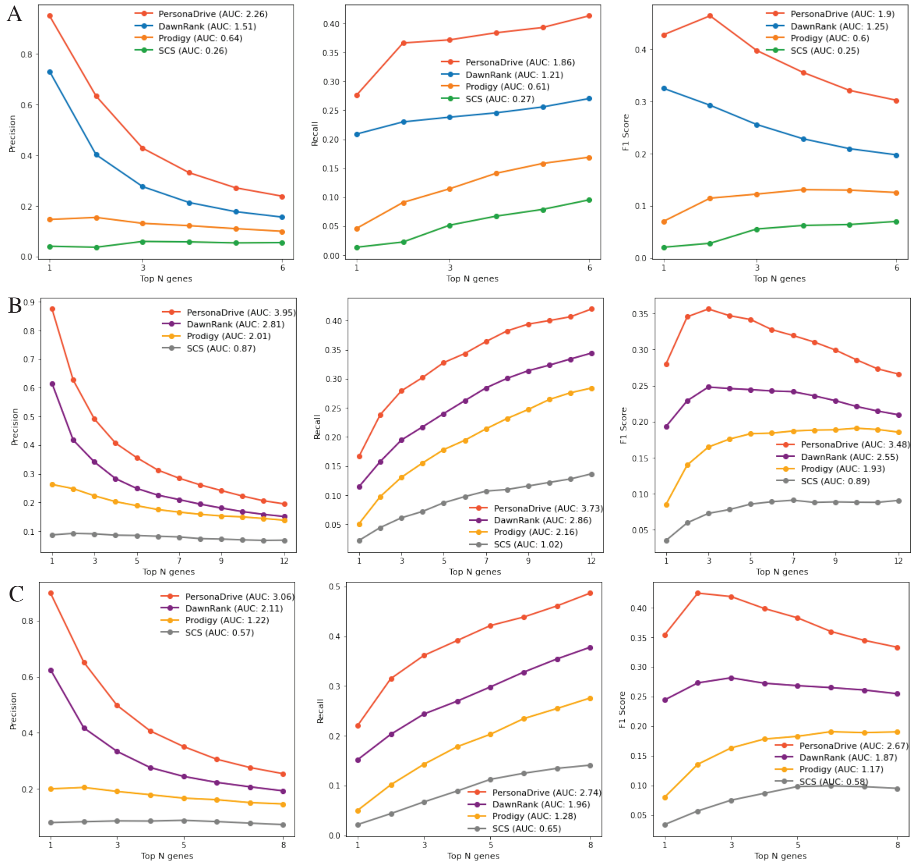


Supplementary Figure 9: Comparison of the PersonaDrive outputs with those of the three alternative methods, DawnRank, SCS, and PRODIGY in terms of average precision, recall, and F1 values for TCGA LUAD dataset where STRING network is used as the input interaction network. A) $CGC_{specific}$ genes are used as reference set. B) $NCG_{CGC}$ genes are used as reference set. C) $CancerMine_{CGC}$ genes are used as reference set.

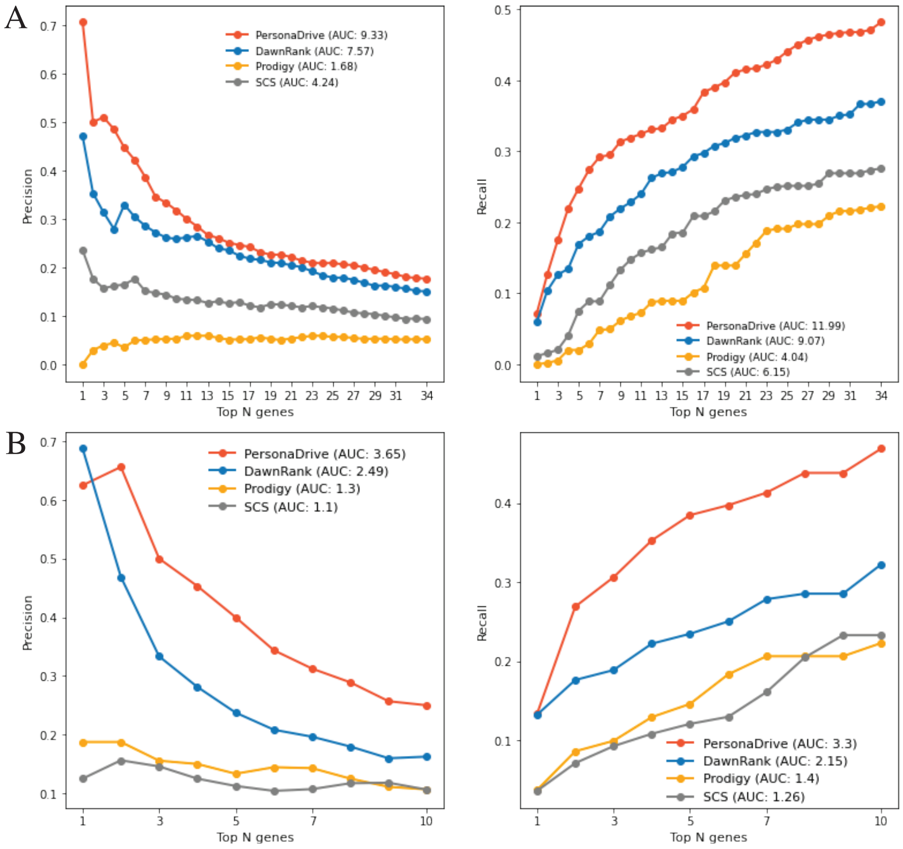


Supplementary Figure 10: Comparison of the PersonaDrive outputs with those of the three alternative methods, DawnRank, SCS, and PRODIGY in terms of average precision, and recall values for A) CCLE COAD cell lines and B) CCLE LUAD cell lines. DawnRank network is used as the input interaction network and the reference set is defined for each cell line based on the targets of sensitive drugs.

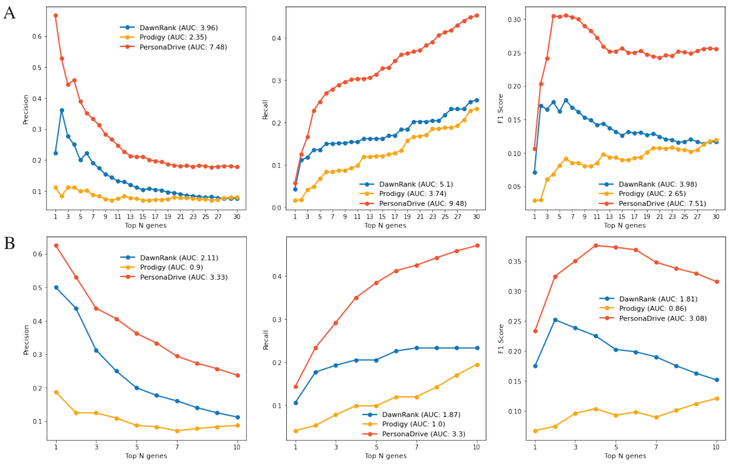


Supplementary Figure 11: Comparison of the PersonaDrive outputs with those of the three alternative methods, DawnRank, SCS, and PRODIGY in terms of average precision, recall, and F1 values for A) CCLE COAD cell lines and B) CCLE LUAD cell lines. STRING network is used as the input interaction network and the reference set is defined for each cell line based on the targets of sensitive drugs.

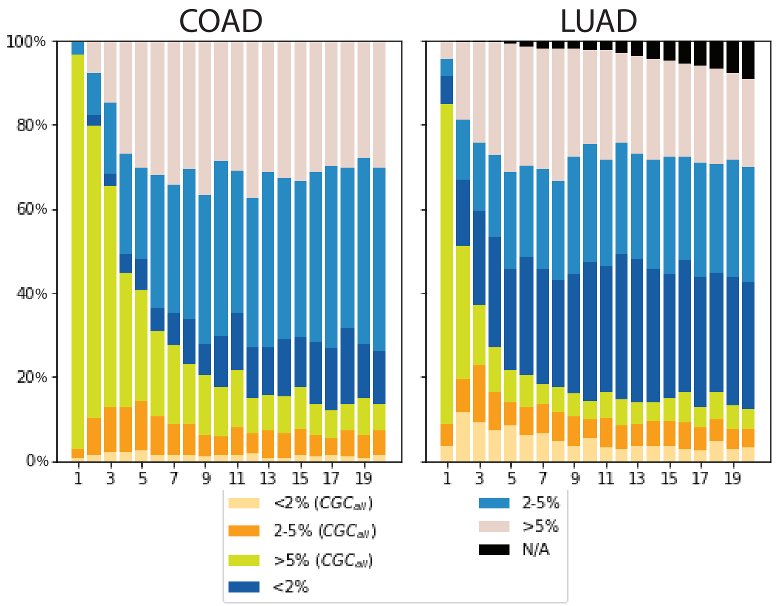


Supplementary Figure 12: Analysis of top 20 ranked genes by PersonaDrive. The figure demonstrates the fraction of samples for which the $k^{th}$ gene belongs to the respective frequency bin. N/A denotes the number of samples for which PersonaDrive outputs a ranked list of less than $k$ mutated genes.

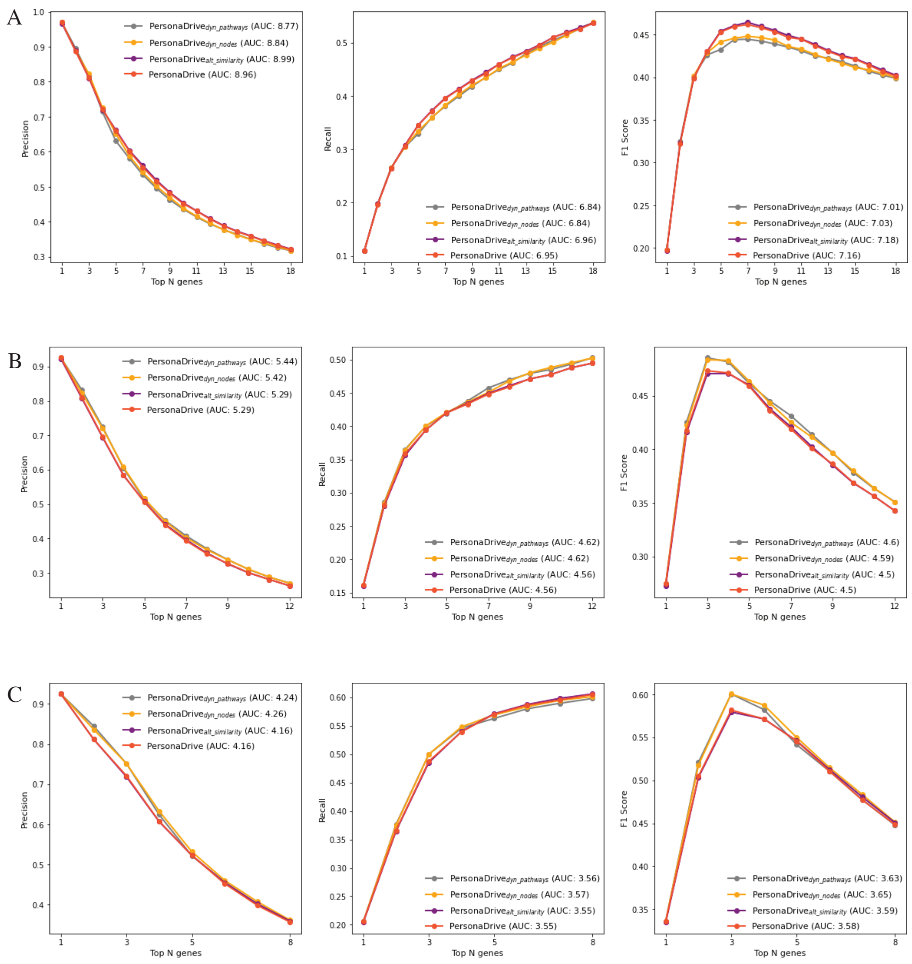


Supplementary Figure 13: Comparison of PersonaDrive with three alternative versions in terms of average precision, recall, and F1 values for TCGA COAD dataset where DawnRank network is used as the input interaction network. A) $CGC_{all}$ genes are used as reference, B) $NCG_{all}$ genes are used as reference, C) $CancerMine_{all}$ genes are used as reference.

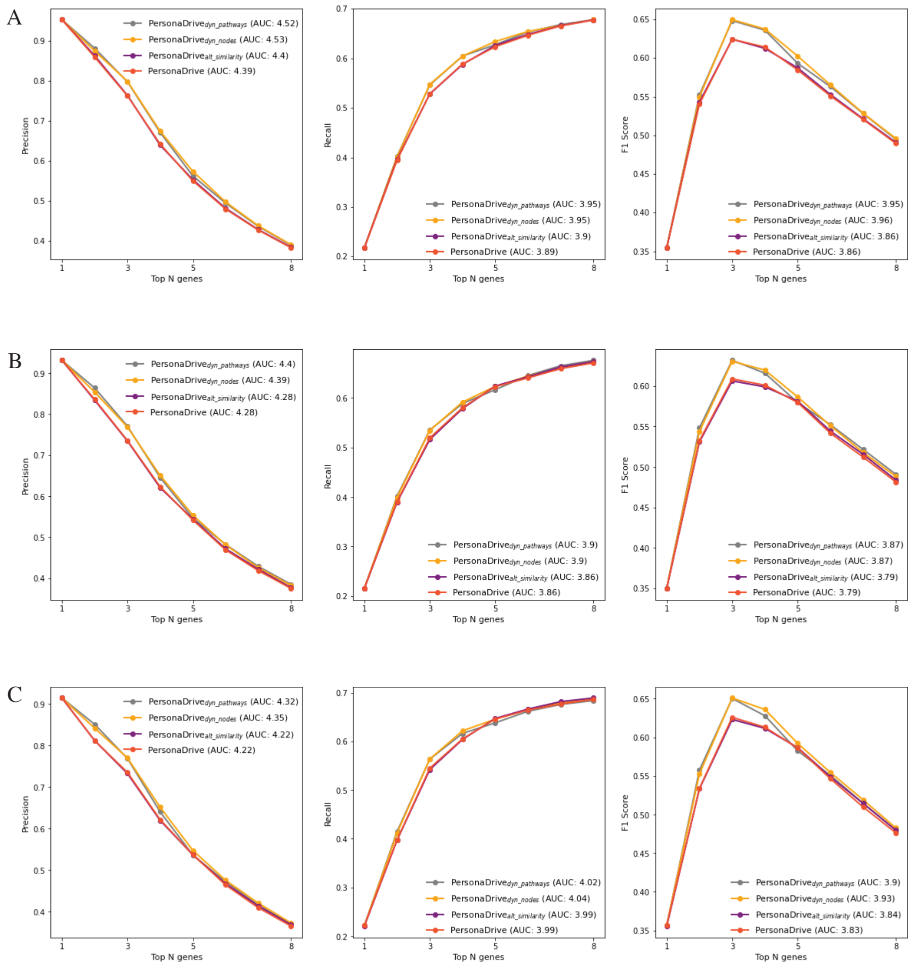


Supplementary Figure 14: Comparison of PersonaDrive with three alternative versions in terms of average precision, recall, and F1 values for TCGA COAD dataset where DawnRank network is used as the input interaction network. A) $CGC_{specific}$ genes are used as reference, B) $NCG_{CGC}$ genes are used as reference, C) $CancerMine_{CGC}$ genes are used as reference.

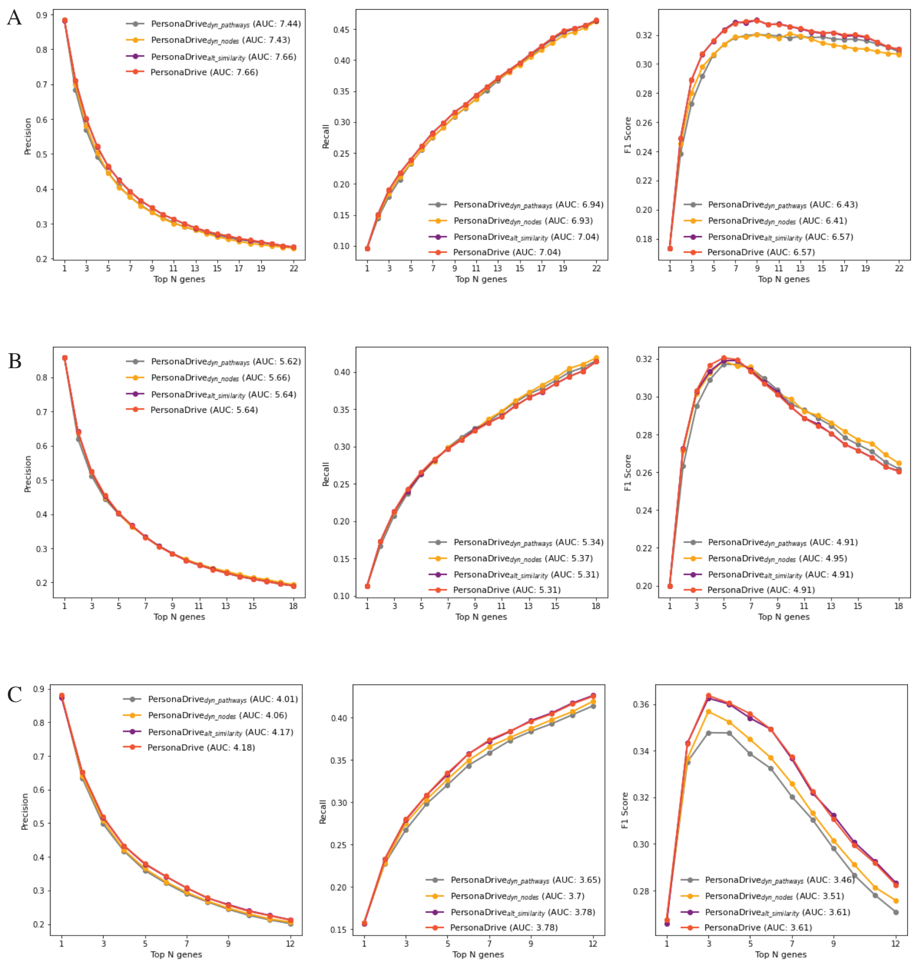


Supplementary Figure 15: Comparison of PersonaDrive with three alternative versions in terms of average precision, recall, and F1 values for TCGA LUAD dataset where DawnRank network is used as the input interaction network. A) $CGC_{all}$ genes are used as reference, B) $NCG_{all}$ genes are used as reference, C) $CancerMine_{all}$ genes are used as reference.

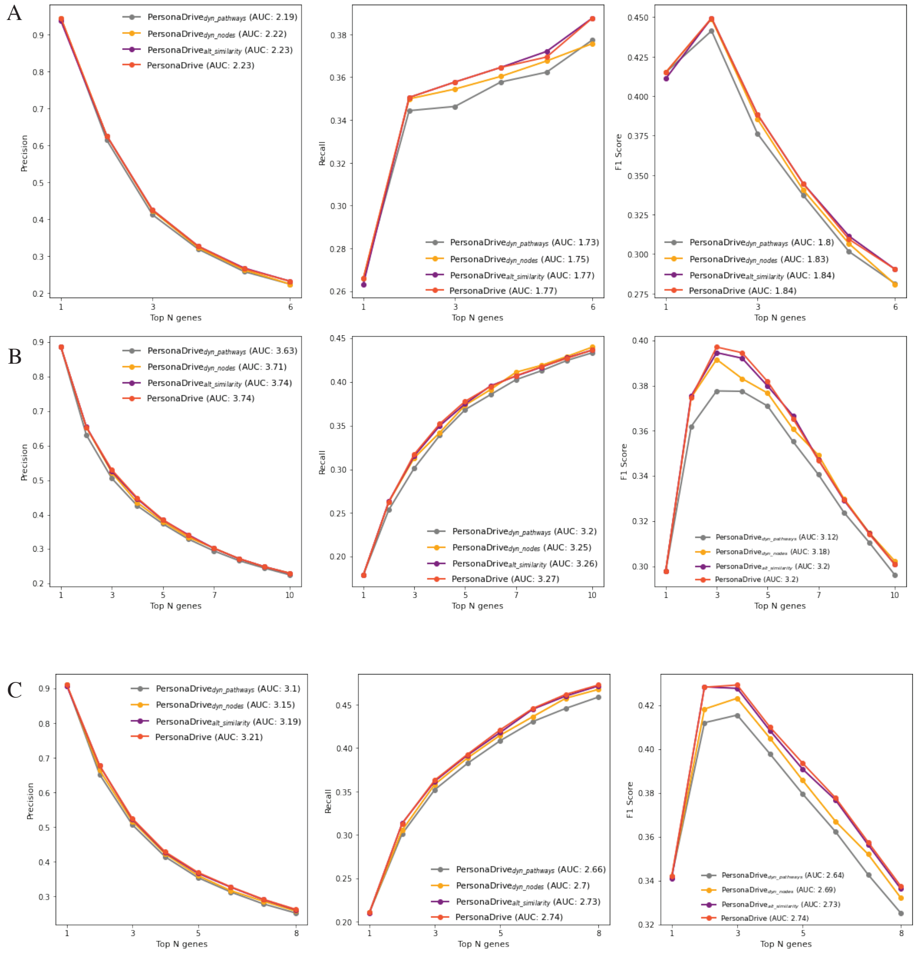


Supplementary Figure 16: Comparison of PersonaDrive with three alternative versions in terms of average precision, recall, and F1 values for TCGA LUAD dataset where DawnRank network is used as the input interaction network. A) $CGC_{specific}$ genes are used as reference, B) $NCG_{CGC}$ genes are used as reference, C) $CancerMine_{CGC}$ genes are used as reference.

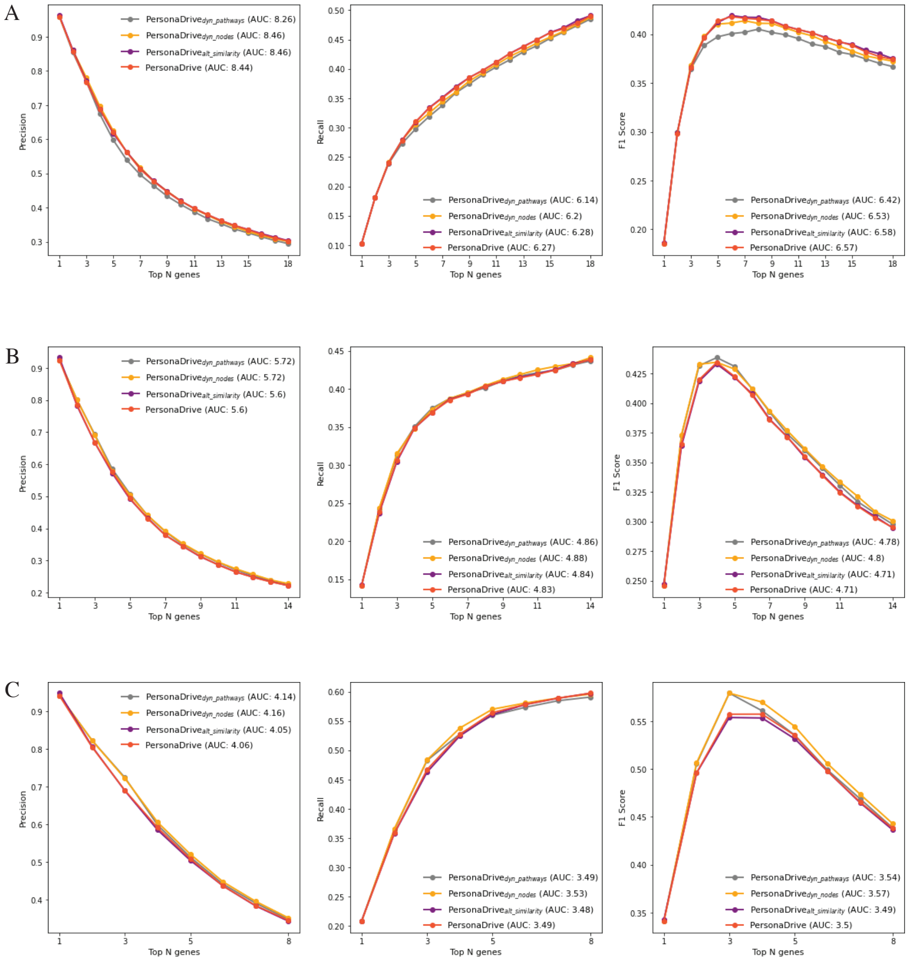


Supplementary Figure 17: Comparison of PersonaDrive with three alternative versions in terms of average precision, recall, and F1 values for TCGA COAD dataset where STRING network is used as the input interaction network. A) $CGC_{all}$ genes are used as reference, B) $NCG_{all}$ genes are used as reference, C) $CancerMine_{all}$ genes are used as reference.

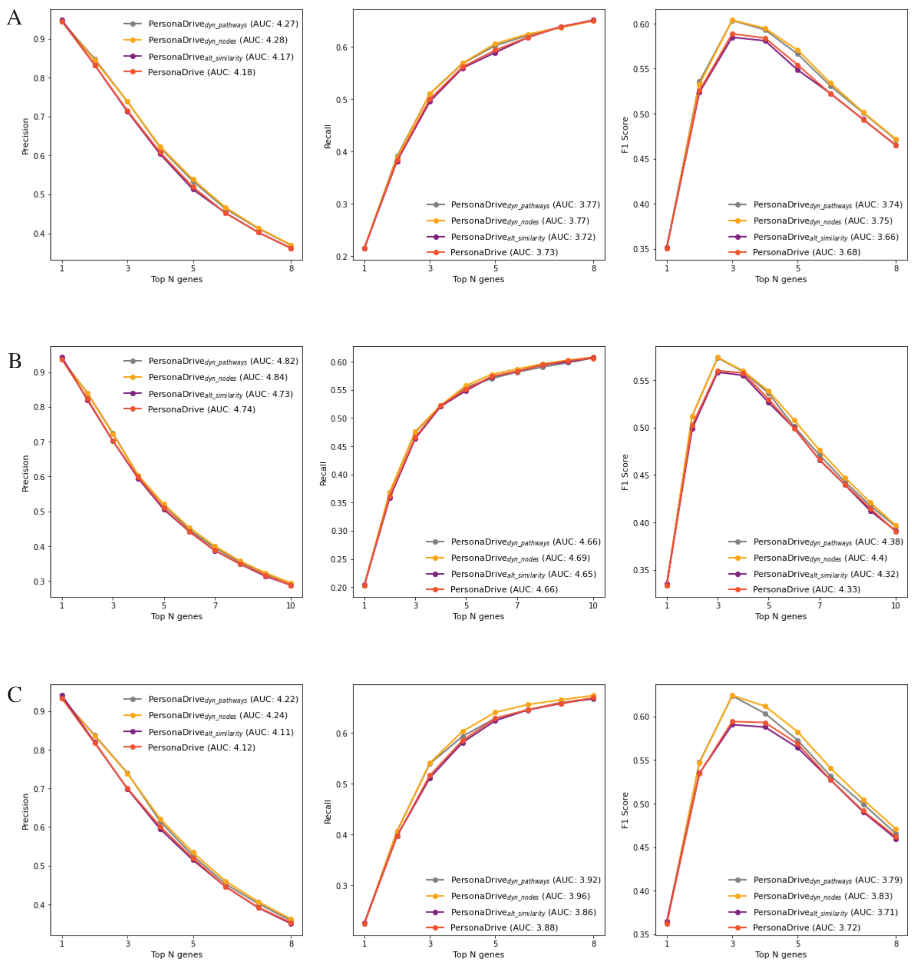


Supplementary Figure 18: Comparison of PersonaDrive with three alternative versions in terms of average precision, recall, and F1 values for TCGA COAD dataset where STRING network is used as the input interaction network. A) $CGC_{specific}$ genes are used as reference, B) $NCG_{CGC}$ genes are used as reference, C) $CancerMine_{CGC}$ genes are used as reference.

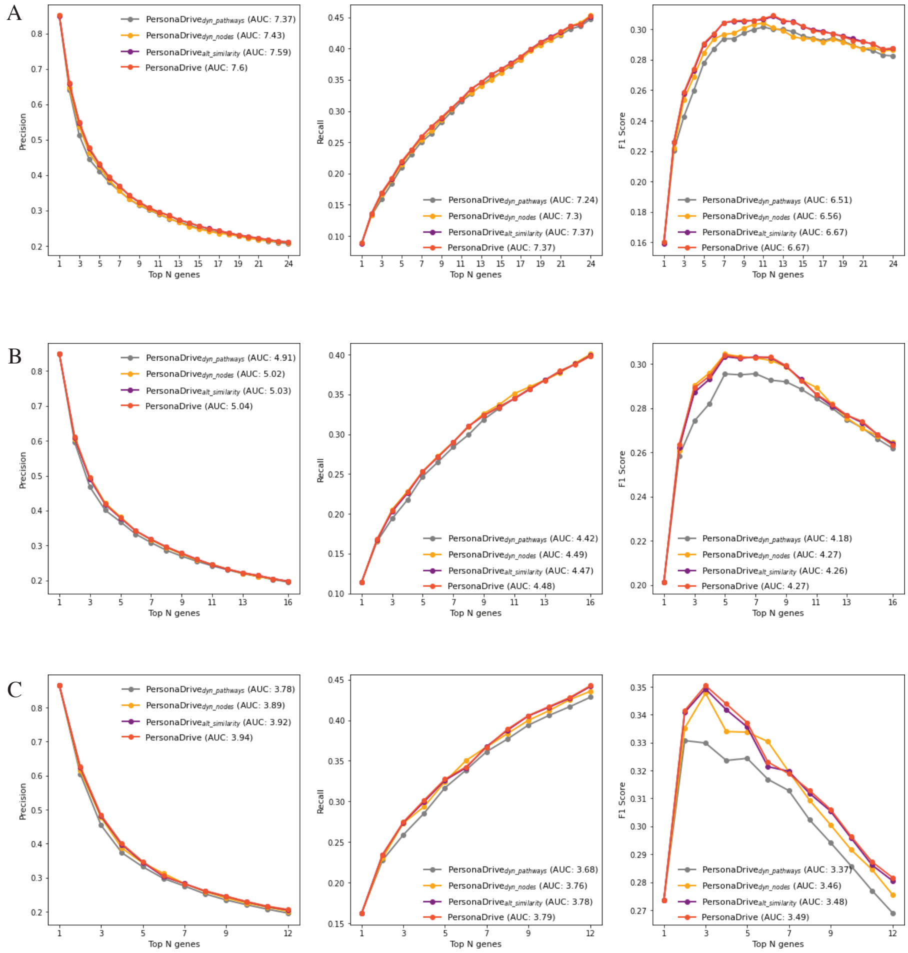


Supplementary Figure 19: Comparison of PersonaDrive with three alternative versions in terms of average precision, recall, and F1 values for TCGA LUAD dataset where STRING network is used as the input interaction network. A) $CGC_{all}$ genes are used as reference, B) $NCG_{all}$ genes are used as reference, C) $CancerMine_{all}$ genes are used as reference.

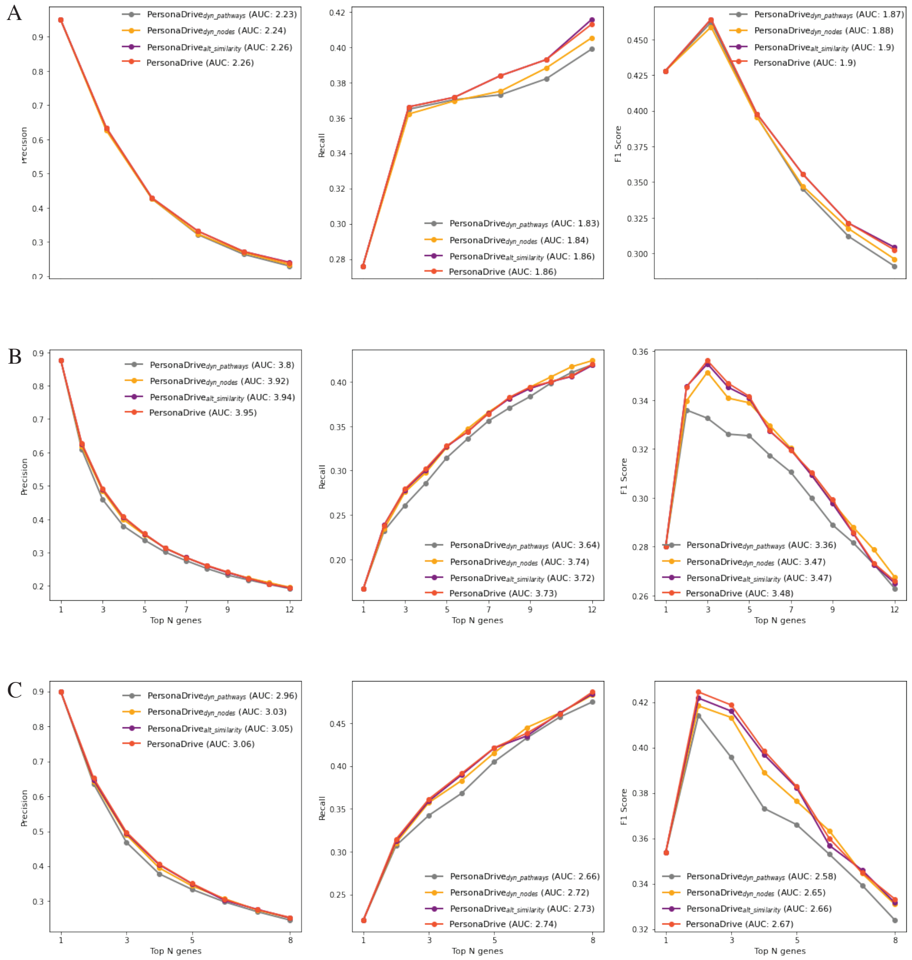


Supplementary Figure 20: Comparison of PersonaDrive with three alternative versions in terms of average precision, recall, and F1 values for TCGA LUAD dataset where STRING network is used as the input interaction network. A) $CGC_{specific}$ genes are used as reference, B) $NCG_{CGC}$ genes are used as reference, C) $CancerMine_{CGC}$ genes are used as reference.

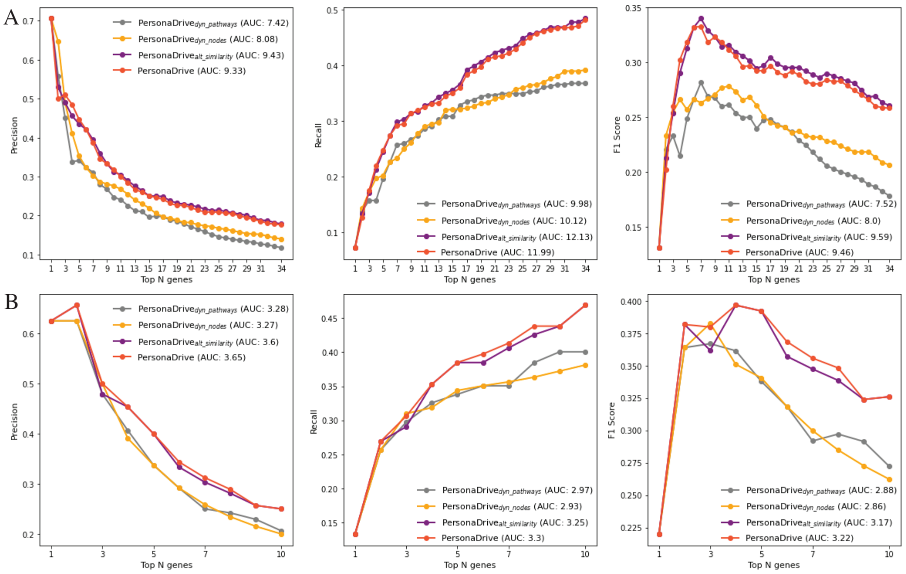


Supplementary Figure 21: Comparison of PersonaDrive with three alternative versions in terms of average precision, recall, and F1 values for A) CCLE COAD cell lines and B) CCLE LUAD cell lines. DawnRank network is used as the input interaction network and the reference set is defined for each cell line based on the targets of sensitive drugs.

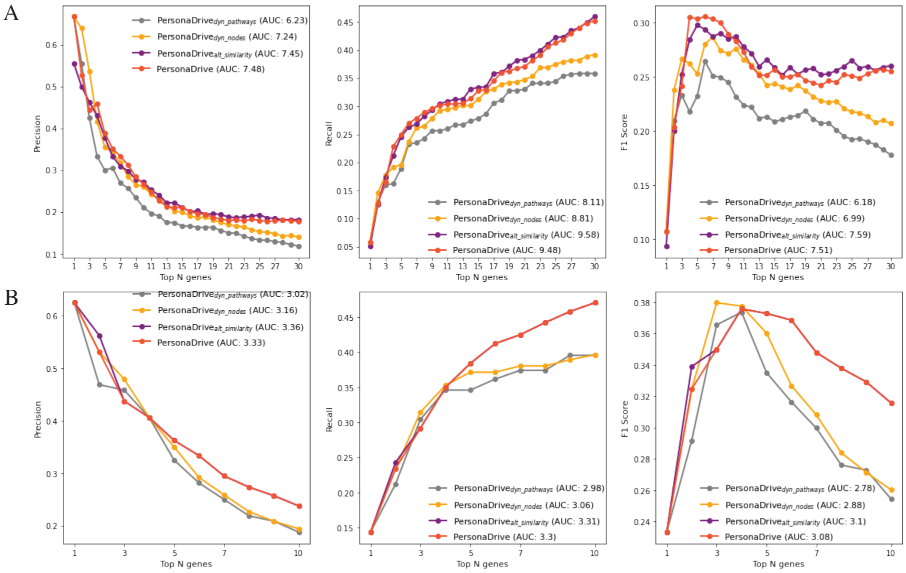


Supplementary Figure 22: Comparison of PersonaDrive with three alternative versions in terms of average precision, recall, and F1 values for A) CCLE COAD cell lines and B) CCLE LUAD cell lines. STRING network is used as the input interaction network and the reference set is defined for each cell line based on the targets of sensitive drugs.

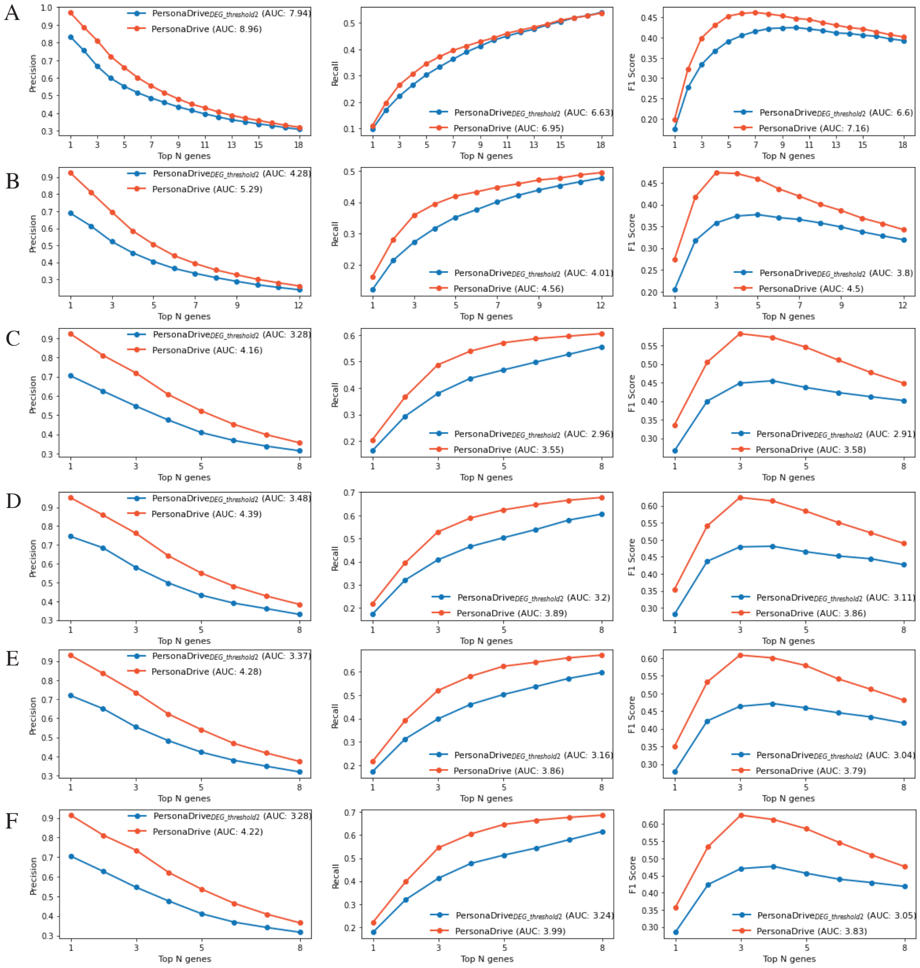


Supplementary Figure 23: Comparison of PersonaDrive with respect to the z-score threshold employed in determining DEGs. The comparison is in terms of average precision, recall, and F1 values for TCGA COAD dataset where DawnRank network is used as the input interaction network. A) $CGC_{all}$ genes are used as reference, B) $NCG_{all}$ genes are used as reference, C) $CancerMine_{all}$ genes are used as reference, D) $CGC_{specific}$ genes are used as reference, E) $NCG_{CGC}$ genes are used as reference, F) $CancerMine_{CGC}$ genes are used as reference.

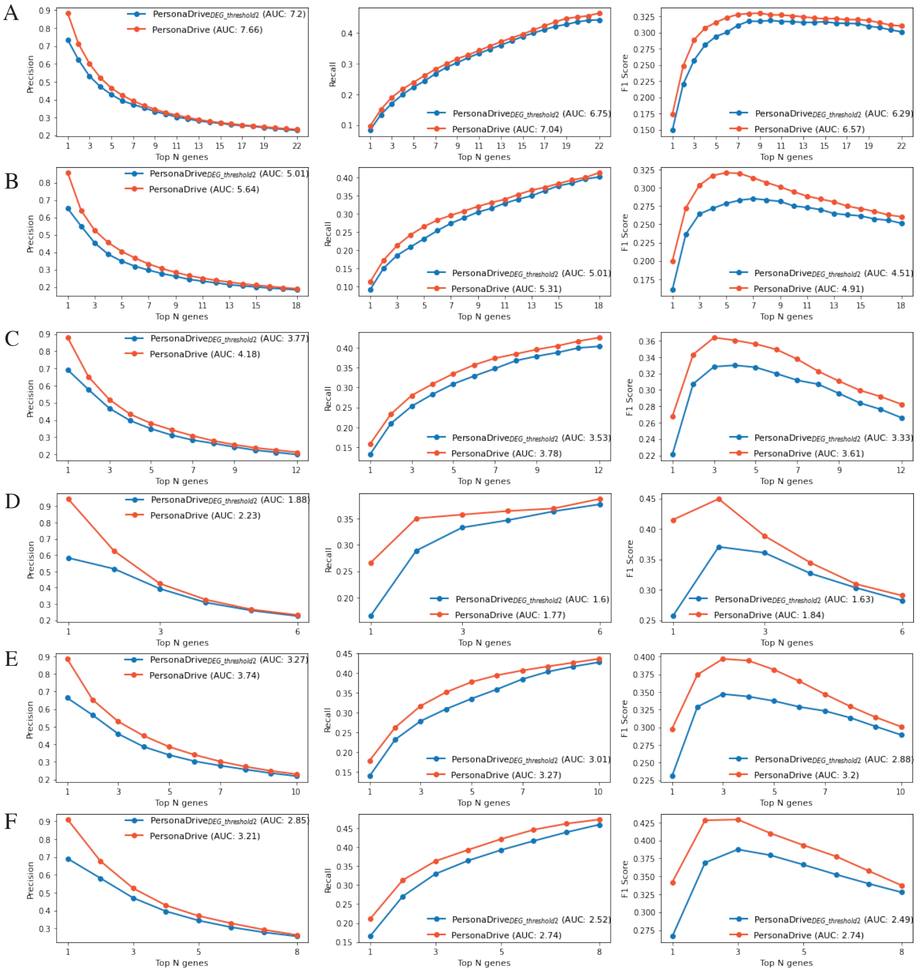


Supplementary Figure 24: Comparison of PersonaDrive with respect to the z-score threshold employed in determining DEGs. The comparison is in terms of average precision, recall, and F1 values for TCGA LUAD dataset where DawnRank network is used as the input interaction network. A) $CGC_{all}$ genes are used as reference, B) $NCG_{all}$ genes are used as reference, C) $CancerMine_{all}$ genes are used as reference, D) $CGC_{specific}$ genes are used as reference, E) $NCG_{CGC}$ genes are used as reference, F) $CancerMine_{CGC}$ genes are used as reference.

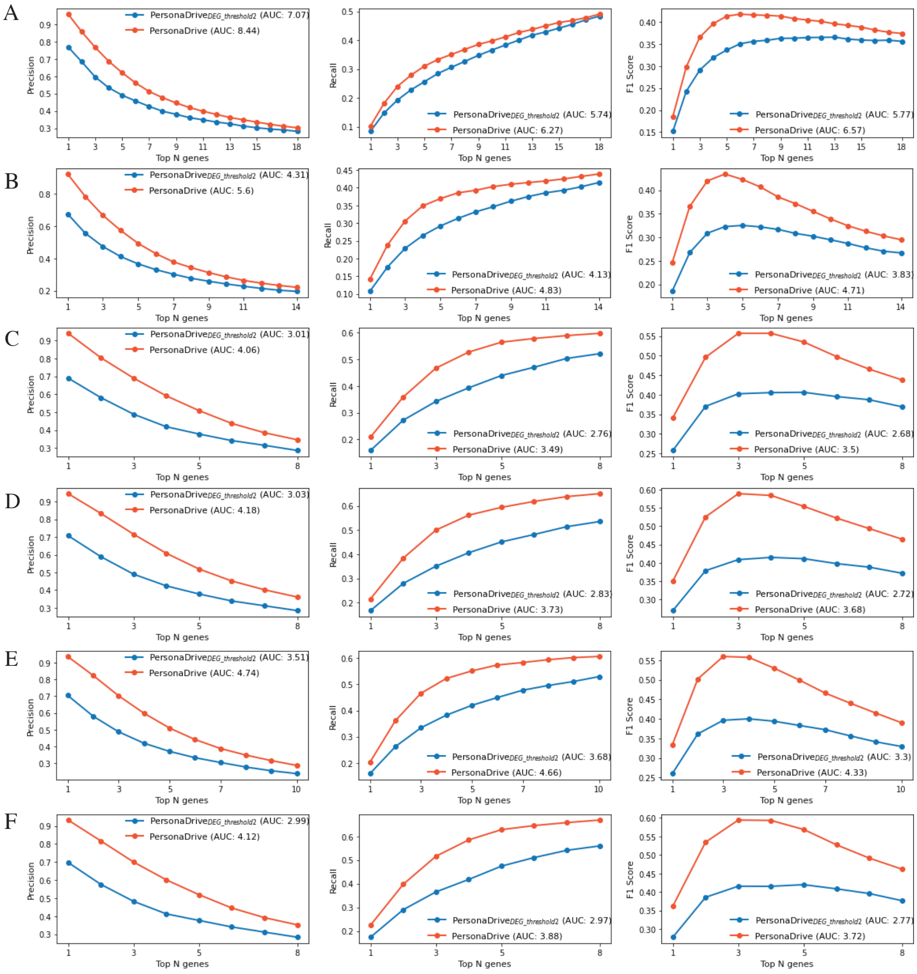


Supplementary Figure 25: Comparison of PersonaDrive with respect to the z-score threshold employed in determining DEGs. The comparison is in terms of average precision, recall, and F1 values for TCGA COAD dataset where STRING network is used as the input interaction network. A) $CGC_{all}$ genes are used as reference, B) $NCG_{all}$ genes are used as reference, C) $CancerMine_{all}$ genes are used as reference, D) $CGC_{specific}$ genes are used as reference, E) $NCG_{CGC}$ genes are used as reference, F) $CancerMine_{CGC}$ genes are used as reference.

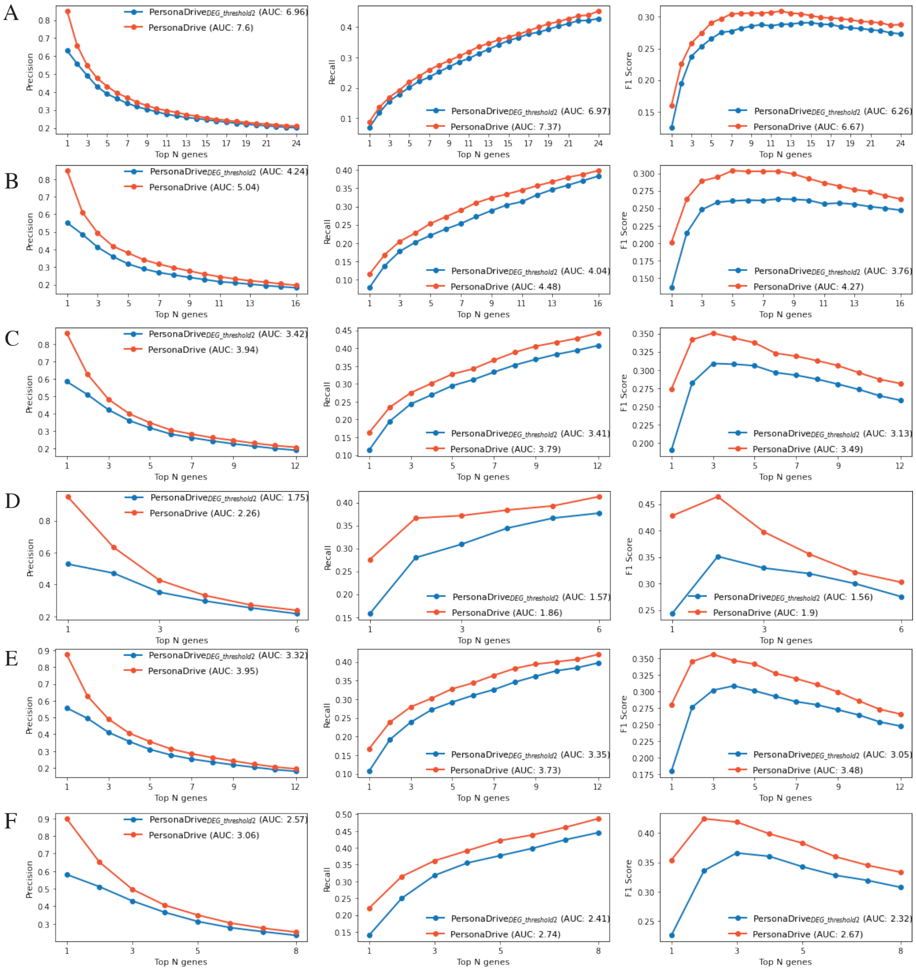


Supplementary Figure 26: Comparison of PersonaDrive with respect to the z-score threshold employed in determining DEGs. The comparison is in terms of average precision, recall, and F1 values for TCGA LUAD dataset where STRING network is used as the input interaction network. A) $CGC_{all}$ genes are used as reference, B) $NCG_{all}$ genes are used as reference, C) $CancerMine_{all}$ genes are used as reference, D) $CGC_{specific}$ genes are used as reference, E) $NCG_{CGC}$ genes are used as reference, F) $CancerMine_{CGC}$ genes are used as reference.

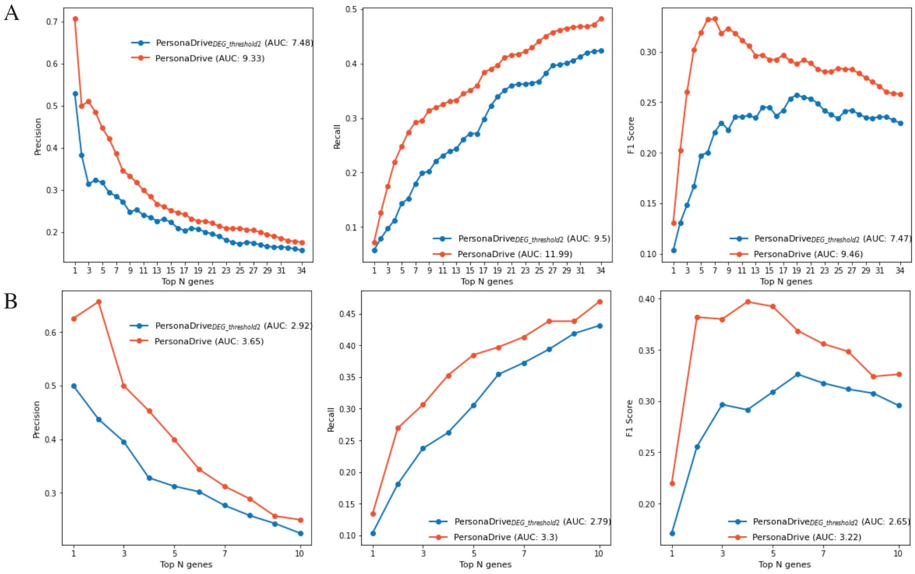

Supplementary Figure 27: Comparison of PersonaDrive with respect to the z-score threshold employed in determining DEGs. The comparison is in terms of average precision, recall, and F1 values for A) CCLE COAD cell lines and B) CCLE LUAD cell lines. DawnRank network is used as the input interaction network and the reference set is defined for each cell line based on the targets of sensitive drugs.

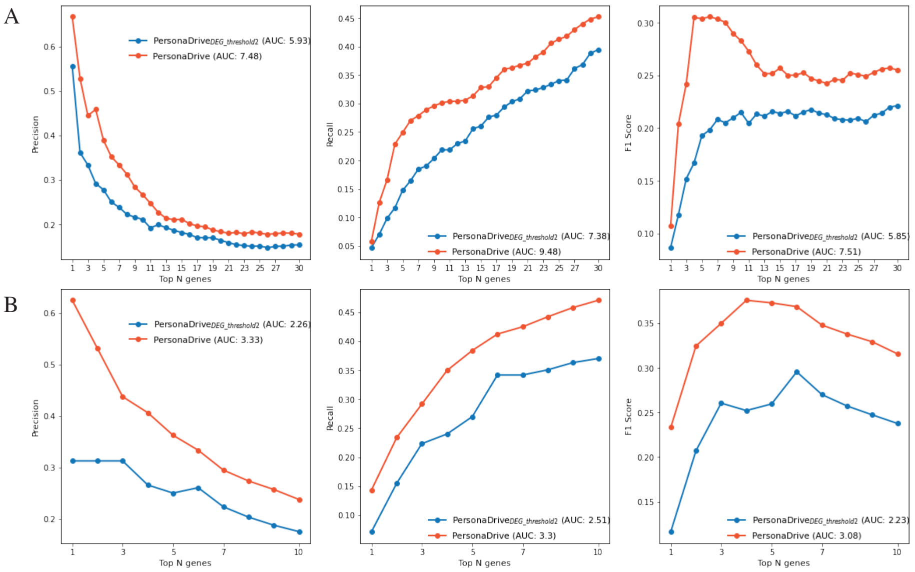


Supplementary Figure 28: Comparison of PersonaDrive with respect to the z-score threshold employed in determining DEGs. The comparison is in terms of average precision, recall, and F1 values for A) CCLE COAD cell lines and B) CCLE LUAD cell lines. STRING network is used as the input interaction network and the reference set is defined for each cell line based on the targets of sensitive drugs.

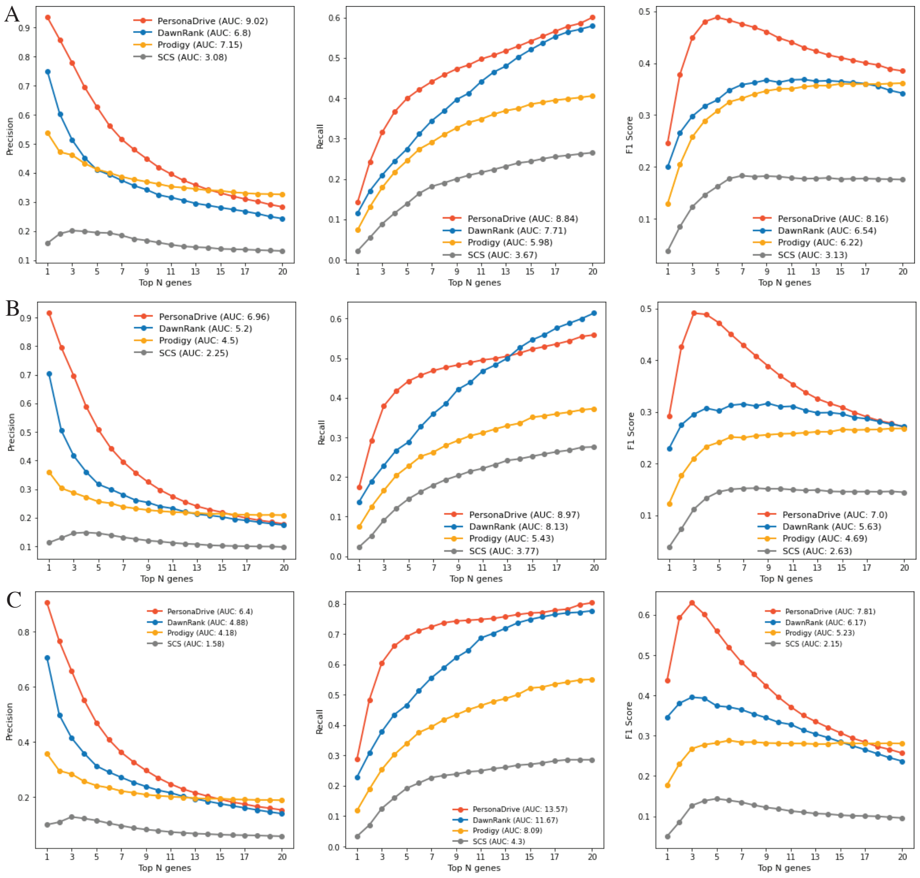


Supplementary Figure 29: Comparison of PersonaDrive with three alternative methods in terms of average precision, recall, and F1 values for TCGA COAD dataset retrieved from [2] where STRING network is used as the input interaction network. In these evaluations we employ the unmodified REA strategy which is the original evaluation strategy proposed in [2]. A) $CGC_{all}$ genes are used as reference. B) $NCG_{all}$ genes are used as reference. C) $CancerMine_{all}$ genes are used as reference.

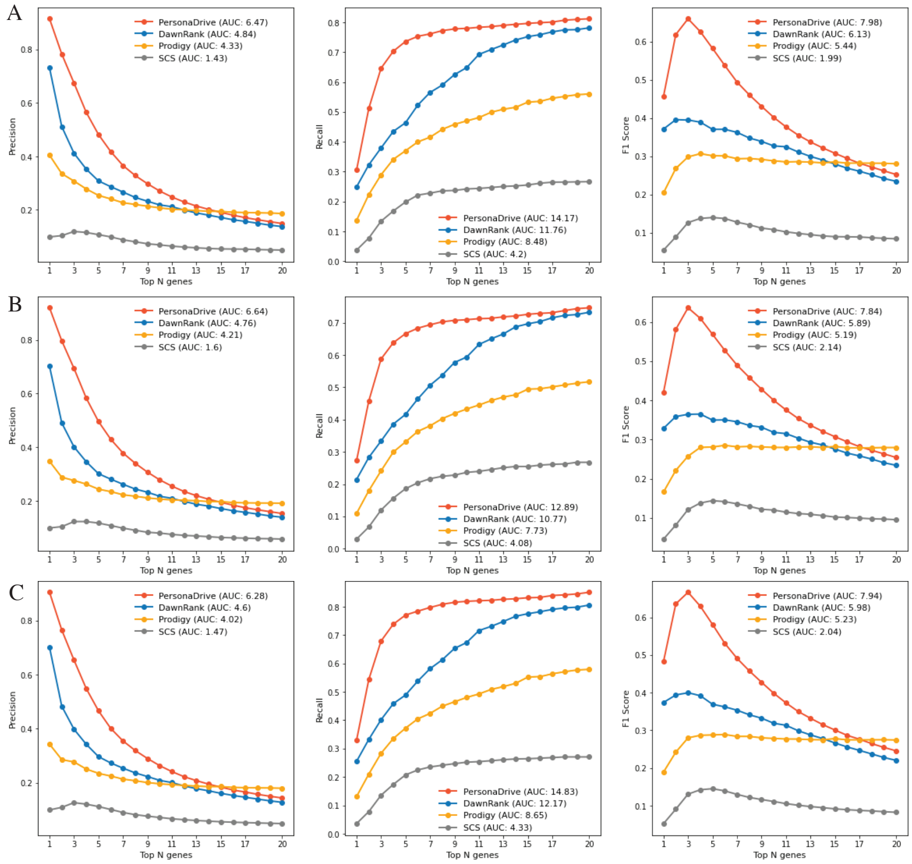


Supplementary Figure 30: Comparison of PersonaDrive with three alternative methods in terms of average precision, recall, and F1 values for TCGA COAD dataset retrieved from [2] where STRING network is used as the input interaction network. In these evaluations we employ the unmodified REA strategy which is the original evaluation strategy proposed in [2]. A) $CGC_{specific}$ genes are used as reference. B) $NCG_{CGC}$ genes are used as reference. C) $CancerMine_{CGC}$ genes are used as reference.

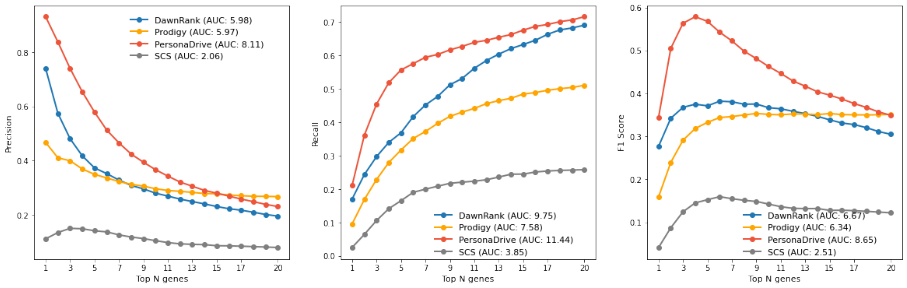


Supplementary Figure 31: Comparison of PersonaDrive with three alternative methods in terms of average precision, recall, and F1 values for TCGA COAD dataset. Both the cancer data and the *CGC* reference set are retrieved from [2]. Here, STRING network is used as the input interaction network. In these evaluations we employ the unmodified REA strategy which is the original evaluation strategy proposed in [2].
